# AAV VP1 unique region (VP1u) determines GPR108 dependence for AAV transduction of human airway epithelium and its rescue by Doxorubicin

**DOI:** 10.64898/2026.07.10.737643

**Authors:** Siyuan Hao, Ariful Habib, Xiujuan Zhang, Kang Ning, Soo Yuen Park, Shane Mcfarlin, Cagla Aksu Kuz, Donovan Richart, Fang Cheng, Ziying Yan, Jianming Qiu

## Abstract

rAAV2.5T was identified through directed evolution of an AAV capsid library in polarized human airway epithelium (HAE) cultured at an air-liquid interface (ALI). The capsid gene of rAAV2.5T is a chimera of the N-terminal unique region of AAV2 VP1 (VP1u) and the VP2 and VP3 regions of AAV5 with a single A581T substitution at the variable region (VR) VIII of the capsids. GPR108, a G protein-coupled receptor, is known as an essential host factor for the transduction of rAAV2 but not of rAAV5. Both AAV2 and AAV5 VP1u colocalized well with GPR108 and, to a lesser extent, with the *trans*-Golgi network (TGN). GPR108 knockout (KO) abolished rAAV2.5T transduction in both HeLa cells and HAE-ALI cultures. Remarkably, short-term treatment with doxorubicin (DOX) at 2 µM completely restored transduction, indicating that DOX can compensate for the loss of GPR108 function. DOX enhanced rAAV2.5T transduction by 50-100-fold in wild-type HAE-ALI cultures and by over 300-fold in the GPR108-deficient cultures. Mechanistic studies demonstrated that this enhancement resulted from altered intracellular trafficking that promoted efficient vector nuclear import, rather than increased vector internalization, proteasome inhibition, or activation of the DNA damage response. Importantly, we identified that the N-terminal 15 amino acids of AAV2 VP1u as the primary determinant of rAAV2.5T dependence on GPR108 for transduction. Collectively, these findings demonstrate that productive transduction of rAAV2.5T in polarized HAE cultures depends on GPR108-mediated intracellular trafficking that limits efficient nuclear entry, and that DOX can relieve this constraint by promoting efficient vector import.

**Significance:** AAV2.5T is an airway-tropic vector with considerable promise for pulmonary gene therapy. We found that host factor GPR108 is required for rAAV2.5T trafficking from the TGN to the nucleus and that this step constitutes a major bottleneck to productive transduction in polarized HAE. In contrast, KIAA0319L (AAVR) plays a key role in AAV intracellular trafficking from the endosome to the TGN but not in internalization into polarized HAE during apical transduction. Transient treatment with low-dose doxorubicin (DOX, 2 µM) enhanced rAAV2.5T transduction in HAE by 50-100-fold through a significant increase in vector nuclear import. Notably, DOX can overcome the transduction deficit caused by GPR108 deficiency, but not that caused by AAVR deficiency. Mechanistically, the N-terminal 15 amino acids of the VP1u confer GPR108 dependence during rAAV2.5T apical transduction of polarized HAE. DOX bypasses this requirement by promoting efficient nuclear import without affecting vector internalization, inhibiting proteasomes, or inducing DNA damage response.

## Introduction

Adeno-associated viruses (AAVs), members of the *Dependoparvovirus* subfamily of the *Parvoviridae* family, have a T=1 icosahedral structure that packages a single-stranded (ss)DNA genome of 4.7 kb (2). Recombinant AAVs (rAAVs) have been used in numerous clinical trials to treat a wide variety of monogenic disorders across multiple target organs, including the eye, liver, skeletal muscle, and lung (3–5). To date, the U.S. Food and Drug Administration (FDA) has approved more than nine rAAV-based gene therapy products, including Luxturna, Zolgensma and Itvisma, Roctavian and Hemgenix, Beqvez, Elevidys, Kebilidi, and Otarmeni for the treatment of inherited retinal dystrophy, spinal muscular atrophy, hemophilia A and B, Duchenne muscular dystrophy, aromatic L-amino acid decarboxylase deficiency, and *Otoferlin*-associated hearing loss, respectively, highlighting the clinical success of rAAV vectors as *in vivo* gene delivery platforms (5). Despite the remarkable clinical success of rAAV-based gene therapy, efficient transduction in many tissues still requires relatively high vector doses (6). This limitation is due, in part, to an incomplete understanding of the mechanisms governing rAAV entry, intracellular trafficking, nuclear import, and genome conversion, all of which are required for productive transgene expression (7,8).

AAV capsid, composed of viral proteins VP1, VP2, and VP3 at a stoichiometry of 1:1:10, determines the cell and tissue tropism (9). The active interaction between the capsid and its cellular surface receptor(s), as well as intracellular host factors, remains poorly understood (10). The initial interaction of the AAV capsid with the host is the attachment that is mediated by capsid-specific binding to glycan moieties (attachment receptors) on the cell surface (11–13).

Upon attachment, AAV uses proteinaceous receptors to trigger cellular internalization. KIAA0319L, termed AAVR, is a multiple-serotype proteinaceous receptor of AAVs in Clades A-F and Clade H (AAV5) (14,15), whereas sialomucin CD164 is an essential entry receptor for Clade G AAVs, including AAV4, AAV11, AAV12, and AAVrh32.33 (16).

Following internalization through endocytosis, AAV capsids interact with multiple host factors in the cytoplasm that mediate intracellular trafficking and nuclear transport. One of the key players in this process is AAVR, which cycles between the cell surface and the *trans*-Golgi network (TGN) (14). During trafficking, AAVR binds to AAV capsids and interacts with TBC1D23 to mediate the endosome-to-TGN transport of the capsid-AAV complex (17). This routing through acidic endosomal compartments is thought to be critical for extruding the VP1 unique region (VP1u) from the capsid surface (18,19). VP1u harbors a phospholipase A₂ (PLA_2_) domain essential for membrane vesicle escape (20,21). Upon reaching the nucleus, where the vector uncoats to release the single-stranded (ss) DNA genome, which is subsequently converted into double-stranded (ds)DNA, forms episomal structures, and serves as a template for transgene transcription (21). However, AAV post-TGN trafficking toward nuclear import is a bottleneck of productive transduction, evidenced by the accumulation of vectors in perinuclear regions (22,23), and remains largely unclear. GPR108, a G protein-coupled receptor-like transmembrane protein (24), has been identified as an essential host factor for the transduction of multiple AAV serotypes except for AAV5 (25,26).

AAV2.5T is an airway-tropic capsid variant isolated from direct evolution of an AAV2 and AAV5 capsid shuffling library in HAE-ALI (27). Its VP1 is a chimeric protein (N-terminal aa1-128 from AAV2 VP1u plus aa129-725 from AAV5) carrying an A581T mutation. AAV2.5T confers >100-fold enhanced apical transduction in HAE compared to rAAV2 and rAAV5 (27). Our previous study on AAV2.5T transduction of polarized HAE through a genome-wide screen of a CRISPR/Cas9 gRNA library identified that AAVR plays an important role in intracellular trafficking of rAAV2.5T in HAE, but not for vector internalization (28), arguing that AAVR is not a proteinaceous receptor of the apical surface to mediate AAV2.5T entry of polarized HAE (28,29). This screen also suggested a critical role of GPR108 in rAAV2.5T apical transduction (**Fig. S1**), consistent with the notion that the GPR108-dependence of rAAV transduction is dictated by VP1u. GPR108 is primarily localized to the TGN membrane, where it serves as a functional hub for sorting and processing (25,26). Although GPR108 is not a host cell receptor for AAV internalization, it is speculated to play a role in AAV transport from the TGN to the nucleus (25). However, the exact role of GPR108 during AAV trafficking remains unknown.

Despite efficient apical membrane uptake, rAAV2 and rAAV5 transduce polarized HAE inefficiently and require very high vector doses to achieve detectable gene expression (22,23,30,31), indicating a substantial post-entry block to productive transduction. Tripeptidyl proteasome inhibitor LLnL (N-acetyl-L-leucyl-L-leucyl-L-norleucine) or ZLL (carbobenzoxy-L-leucyl-L-leucyl-leucinal) enhanced apical transduction of rAAV2 in polarized HAE (23), and prevented proteasome degradation of AAV capsids in HeLa cells, demonstrating increased levels of ubiquitinated AAV2 and AAV5 capsids (32). Combined administration of anthracycline DOX and LLnL or ZLL synergistically augments rAAV2 and rAAV5 transduction from the apical membrane of polarized HAE-ALI (1). Although AAV2.5T confers >100-fold enhanced apical transduction in HAE compared to rAAV2 and rAAV5 (27), it remains subject to post-entry barriers and requires a high vector dose for efficient transduction. Treatment of DOX alone is sufficient to augment apical transduction of rAAV2.5T in both HAE and ferret airway epithelium (FAE), as well as in ferret lungs (28,33–35). DOX is an FDA-approved pleiotropic anticancer drug (36–38); it exhibits dose-dependent proteasome inhibitory activity (39). Previous studies suggested that DOX itself might also modulate the ubiquitin-proteasome system, as combined use of DOX and LLnL or ZLL synergistically enhanced rAAV2 or rAAV5 transduction in HAE-ALI, and that co-application of DOX and LLnL enhanced rAAVs’ nuclear import (1,40). However, the precise mechanism underlying the DOX-enhanced rAAV2.5T transduction in polarized HAE remains elusive.

In this study, using the polarized HAE-ALI model, we investigated the subcellular localization of GPR108 relative to the TGN and AAV VP1u, and mapped the VP1u domain required for GPR108-dependent transduction of rAAV2.5T. We found that administration of DOX at 2 µM during a short transduction period (∼16 h/overnight) augmented AAV2.5T transduction by 50-100-fold without inhibiting proteasome activity, activation of the DNA damage response (DDR), or causing detectable host DNA damage. Notably, DOX treatment compensated for the loss of GPR108 in GPR108 knockout (KO) HAE-ALI cultures, but did not rescue transduction in AAVR deficient cultures.

## Results

### GPR108 is required for rAAV2.5T transduction but not for rAAV5 transduction

In order to confirm the role of GPR108 in rAAV2.5T transduction, we generated GPR108 KO HeLa (HeLa^GPR-KO^) cells (**Fig S2**). We then transduced both nontarget (NT) control HeLa cells (HeLa^NT^) and HeLa^GPR-KO^ with rAAV2.5T.fLuc-mCherry at a multiplicity of infection (MOI) of 20,000 DNase-resistant particles (DRP)/cell, using rAAV5 as a control for GPR108-independent transduction. At 3 days post-transduction (dpi), we assessed both mCherry expression and luciferase activity. For rAAV5, no statistically significant differences in the expression of mCherry and firefly luciferase (fLuc) activity were observed between the HeLa^NT^ and HeLa^GPR-KO^ cells (**Fig. 1A&B**). Conversely, rAAV2.5T transduction resulted in drastically decreased mCherry expression and a >98% reduction in fLuc activity in HeLa^GPR-KO^ cells compared to HeLa^NT^ cells (**Fig. 1C&D**). Together, these data demonstrate that rAAV2.5T transduction in HeLa cells is GPR108-dependent, whereas rAAV5 transduction is not.

**Fig. 1.**
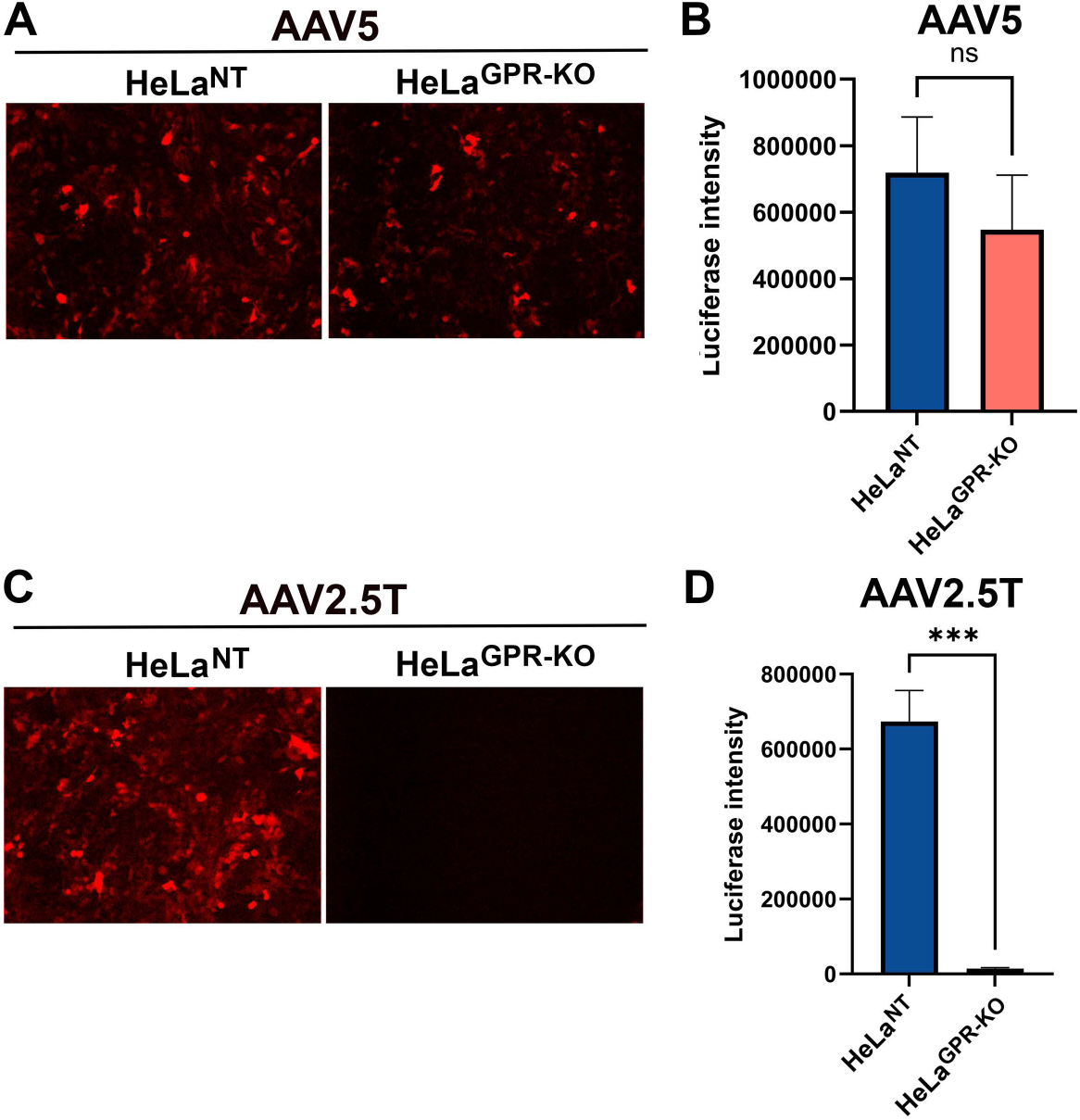
GPR108 is essential for rAAV2.5T but not AAV5 transduction in HeLa cells. rAAV5 (A&B) or rAAV2.5T (C&D) was transduced to non-target (NT) and GPR108-KO HeLa cells, respectively, at an MOI of 20,000 DRP/cell. **(A&C) mCherry expression**. At 72 h post-transduction (hpt), fluorescence was visualized under a NIKON Ti-S fluorescent microscope. Images were taken at a magnification of x 40, and representative images from three independent experiments are shown. **(B&D) Luciferase activity**. At 3 dpt, transduced cells were harvested and analyzed for firefly luciferase activity. Data are presented as the mean ± standard deviation (SD) from three replicates. Statistical significance was determined using an unpaired two-tailed Student’s t-test. ns, not significant; ***, P < 0.001.

While rAAV5 and rAAV2.5T transduced HeLa^NT^ with similar efficiency, the distinct transduction profiles in HeLa^GPR-KO^ cells were likely due to the difference in the VP1u region, as AAV2.5T possesses the AAV2 VP1u (27). To investigate this, we individually expressed Flag-tagged AAV2 VP1u and AAV5 VP1u in HeLa cells and evaluated their colocalization with GPR108. Expressed from plasmid transfection, both AAV2 and AAV5 VP1u were localized predominantly to the nucleus, consistent with the presence of the nuclear localization signal (NLS) at the C-terminus of VP1u (41). Notably, the fractions retained in the cytoplasm were well-colocalized with GPR108 (**Fig. 2A**), and a large portion of each was localized to the TGN, as indicated by TGN46 staining (**Fig. 2B**). Next, we transduced HeLa cells with rAAV2, rAAV5, and rAAV2.5T, respectively. At 8 hours post-transduction (hpt), GPR108 displayed a diffuse cytoplasmic distribution, and a substantial fraction of the capsids from all three vectors accumulated in the cytoplasm and colocalized with GPR108 (**Fig. 3A**). However, TGN46 was more concentrated in one pole of the cytoplasm where AAV capsids localized (**Fig. 3B**).

**Fig. 2.**
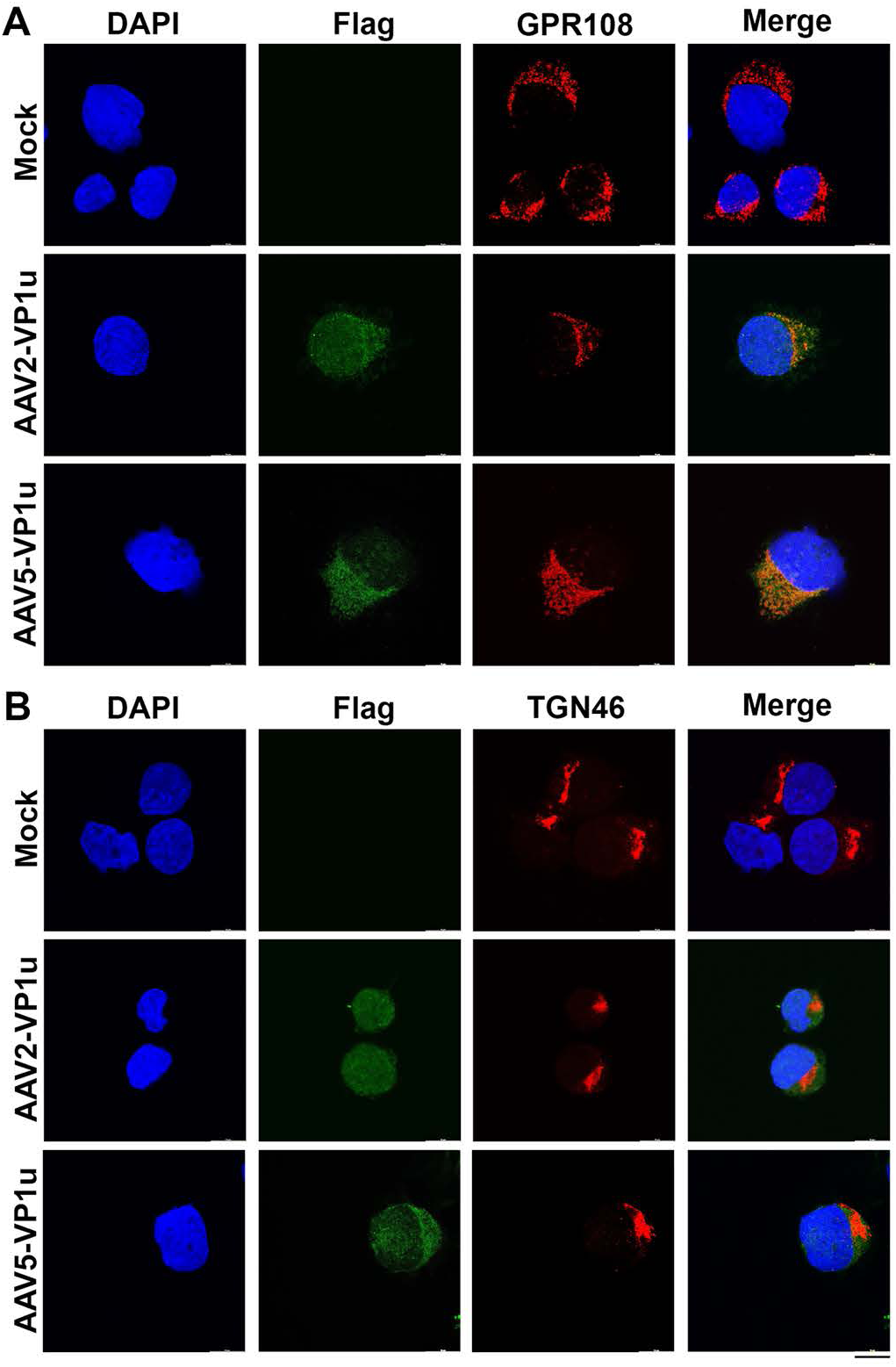
AAV2 and AAV5 VP1u proteins colocalize with GPR108 but show limited association with TGN46 in HeLa cells. HeLa cells were transfected with plasmids expressing Flag-tagged AAV2 VP1u or AAV5 VP1u. At 72 hpt, cells were fixed and stained with anti-Flag and either anti-GPR108 or anti-TGN46 antibodies, as indicated. Confocal immunofluorescence images were acquired using a Leica TCS SP8 STED confocal microscope with a ×100 objective. **(A) VP1 colocalization with GPR108.** Representative images show the intracellular localization and colocalization of AAV2 VP1u^Flag^ or AAV5 VP1u^Flag^ with GPR108. **(B) VP1 colocalization with TGN46.** Representative images show the intracellular localization and association of AAV2 VP1u^Flag^ or AAV5 VP1u^Flag^ with TGN46 (B). Nuclei were stained with DAPI (4′,6-diamidino-2-phenylindole). Scale Bar = 10 µm.

**Fig. 3.**
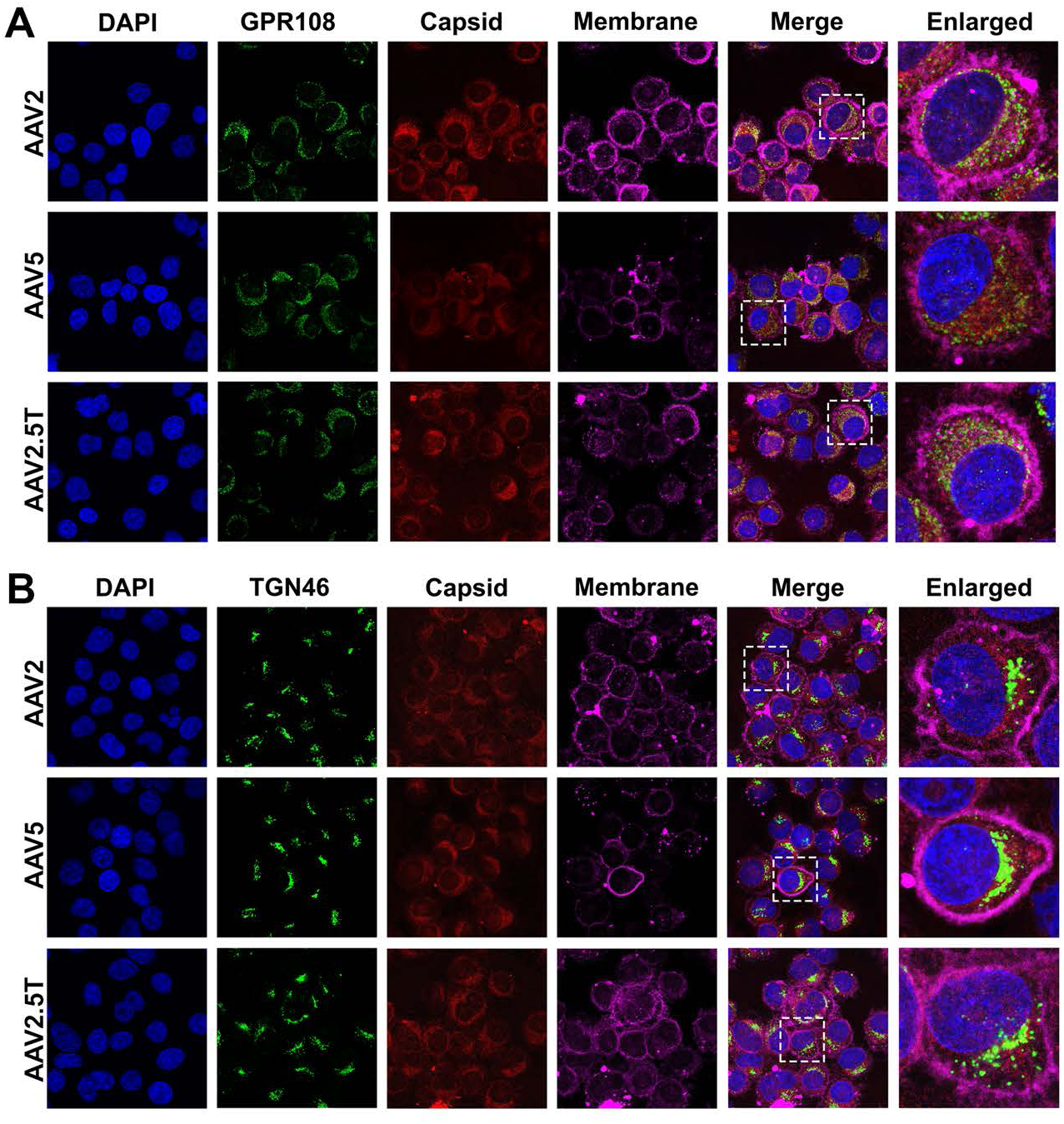
rAAV2, rAAV2.5T, and rAAV5 capsids colocalize with GPR108 and partially associate with TGN46 in HeLa cells. HeLa cells were transduced with rAAV2, rAAV2.5T, or rAAV5 vectors at an MOI of 20,000 DRP/cell. At 8 hpt, the cell membrane was stained with MemBrite Fix 640/660 (Biotium) for 10 min prior to cytospin preparation. Cells were then fixed and immunostained with antibodies against AAV capsids and either GPR108 or TGN46, as indicated. Confocal immunofluorescence images were acquired using a Leica TCS SP8 STED confocal microscope with a ×100 objective. Enlarged views of boxed regions are shown on the right. Representative images show colocalization of rAAV2, rAAV2.5T, and rAAV5 capsids with GPR108- (A) or TGN46-positive compartments (B). Nuclei were stained with DAPI. Cell membrane was stained with MemBrite Fix 640/660 (Magenta).

Taken together, these data suggested that intracellular AAV capsids are present within GPR108-positive compartments during post-entry trafficking. While the capsids traffic to the TGN, their distribution extends beyond the TGN46-positive Golgi structure, suggesting that GPR108 associates with a broader intracellular trafficking network than the TGN alone. Notably, AAV5 capsids were also localized within these GPR108-positive compartments, despite GPR108 not being required for AAV5 transduction.

### The N-terminus of AAV2 VP1u plays a dominant role in conferring both the high transduction efficiency and GPR108 dependence of rAAV2.5T

We next determined the domain in AAV2 VP1u that is responsible for GPR108 dependence. We performed a rational design of rAAV2.5T mutants that have chimeric VP1u of AAV5 and AAV2, as illustrated in **Fig. 4A&B**. AAV5T (C1) had all the AAV5 VP1u, and C2 is a mutant AAV5T capsid that has aa 13-16 of the VP1u replaced with AAV2 VP1u, which remains in the α-helix region of the N-terminus (**Fig. 4C**, C2). C3 to C6 were rAAV5T capsids with increasing lengths of AAV2 VP1u at the N-terminus, which retain structures similar to those of AAV2 and AAV5 VP1u and have 4 α-helices (**Fig. 4C**). C6 has a break after the first α-helix, while the others have a continuous α-helix at the N-terminus.

**Fig. 4.**
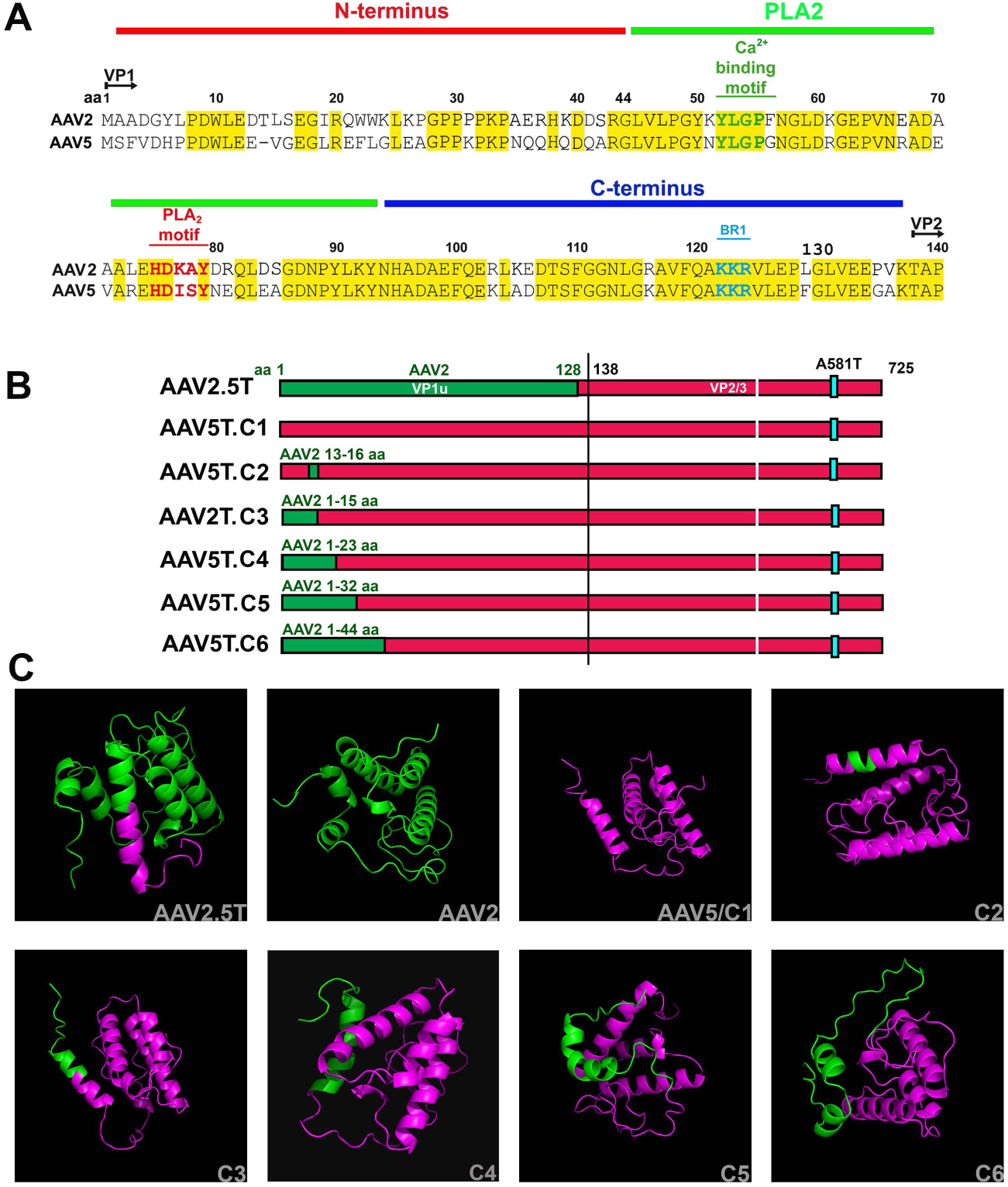
Construction and structural prediction of AAV2.5T, AAV5T, and AAV5T-based VP1u chimeric mutants. **(A) Amino acid sequence alignment of the VP1u of AAV2 and AAV5.** Conserved amino acid residues are highlighted in yellow. Functional regions within VP1u, including the N-terminal region, the phospholipase A2 (PLA_2_) domain including the calcium (Ca^2+^) binding motif and PLA_2_ catalytic motif, and the basic region 1 (BR1) that functions as a nuclear localization signal (NLS), are indicated. **(B) Schematic representation of the VP1u regions of AAV2.5T, AAV5T, and AAV5T-based chimeric mutants (C1–C6).** AAV2-derived VP1u amino acids are shown in green, whereas AAV5-derived sequences are shown in magenta. The cyan box indicates the A581T mutation within the capsid proteins. The vertical lines denote the VP1u/VP2 boundary region. **(C) AlphaFold-predicted structures of AAV2.5T, AAV2, AAV5T, and chimeric VP1u mutants (C2–C6).** AAV2-derived regions are shown in green and AAV5-derived regions in magenta. AAV2.5T and the AAV5T.C6 mutant exhibit a disrupted α-helical structure at the N-terminus of VP1u, whereas AAV5T and mutants AAV5T.C3–C6 retain a more continuous α-helical conformation.

We transduced both HeLa cells and HAE-ALI cultures with AAV2.5T, AAV5T, and VP1u mutants, and measured fLuc activity at 3 dpt. We found that rAAV2.5T transduction was significantly higher than that of rAAV5T and AAV5T.C2-6 mutants in both cultures (**Fig. 5A&B**). Interestingly, in the four transductions of rAAV5T.C3 to C6, we observed a clear trend in which the longer the AAV2 VP1u sequence swapped in the mutants, the higher the transduction efficiency. AAV5T.C6, which has AAV2 VP1u aa 1-44, transduced HeLa cells and HAE-ALI cultures at 74% and 82%, respectively, of the parent AAV2.5T transduction level. To assess whether these VP1u chimeric rAAVs remain PLA_2_ active, we measured PLA_2_ activity on denatured virions. No significant differences were detected between AAV5T and AAV5T.C2-C6, however, all variants exhibited lower PLA_2_ activity compared with AAV2.5T (**Fig. 5E&F**). These results suggested that the N-terminal 44 aa of AAV2 VP1u are largely responsible for the high transduction efficiency of rAAV2.5T in HAE-ALI cultures, and that replacement of the AAV5 VP1u N-terminus with the corresponding AAV2 sequence does not affect VP1u PLA_2_ activity.

**Fig. 5.**
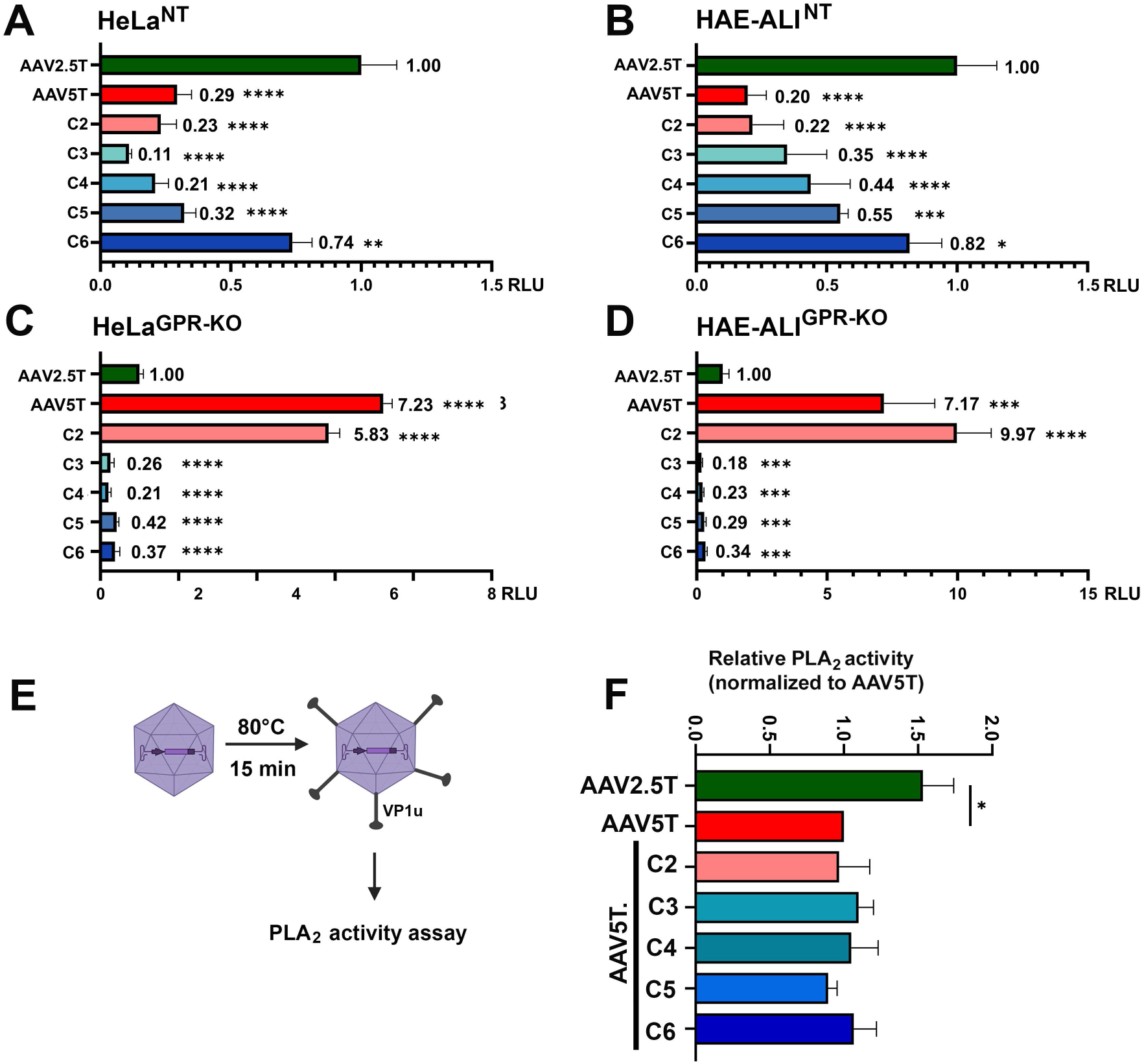
The N-terminus of VP1u determines the GPR108 dependence of rAAV transduction. **(A&B) Transduction efficiencies of AAV2.5T, AAV5T, and AAV5T-based VP1u chimeric mutants in NT control cells.** NT HeLa cells (A) and HAE-ALI cultures (B) were transduced with rAAV vectors at an MOI of 20,000 DRP/cell. Luciferase activity was measured at 5 dpt and normalized to rAAV2.5T transduction, which was set to 1. **(C&D) Transduction efficiencies of AAV2.5T, AAV5T, and VP1u chimeric mutants in GPR108-KO cells.** GPR108-KO HeLa cells (C) and HAE-ALI cultures (D) were transduced with rAAV vectors at an MOI of 20,000 DRP/cell. fLuc activity was measured at 5 dpt and normalized to rAAV2.5T transduction. **(E) Schematic illustration of the phospholipase A2 (PLA_2_) activity assay.** rAAV vectors were heat-treated at 80°C for 15 min to externalize the VP1 unique region (VP1u) prior to measurement of PLA_2_ activity. **(F) Comparison of PLA2 activities among AAV2.5T, AAV5T, and VP1u chimeric mutants (C2–C6).** PLA_2_ activity was measured using a colorimetric assay and normalized to rAAV5T. Data are presented as mean ± SD from three independent experiments. Statistical significance was determined using one-way ANOVA followed by Dunnett’s multiple-comparison test. *, P < 0.05; ***, P < 0.001; ****, P < 0.0001. No significant differences in PLA_2_ activity were observed between AAV5T and mutants C2–C6.

Furthermore, we conducted rAAV mutant transduction in GPR108-KO HeLa cells and HAE-ALI cultures (**Fig. S2A**). We noticed that rAAV2.5T transduction was significantly lower than that of rAAV5T or rAAV5T.C2, and that rAAV5T.C3-6 showed only minimal fLuc activity, >17-fold and 21-fold lower than that of rAAV5T in HeLa^GPR-KO^ and HAE-ALI^GPR-KO^, respectively (**Fig. 5C&D**). AAV5T.C2 that has 4 aa changes from AAV5 ^13^EVG^15^ to AAV2 ^13^DTLS^16^ (**Fig. 4A**) in AAV5 VP1u remained GPR108-independent. However, AAV5T.C3, which possesses the first 15 aa of AAV2 VP1u, has become dependent on GPR108. These data suggest that the minimal N-terminal motif of AAV2 VP1u, aa1–15, plays a critical role in GPR108-dependent transduction.

Taken together, these data suggest that the N-terminus of AAV2 VP1u aa 1-44 confers the high transduction efficiency of rAAV2.5T in HAE-ALI cultures, whereas the N-terminal aa 1-15 region is the major determinant of GPR108-dependent transduction.

### Doxorubicin (DOX) bypasses the requirement for GPR108 during rAAV2.5T transduction

DOX is an FDA-approved chemotherapeutic agent that has also been shown to potently enhance rAAV transduction in polarized HAE and ferret lungs (28,33–35). To evaluate a dose-dependent effect on transducing HAE-ALI and epithelial integrity, we transduced HAE-ALI in the presence of various concentrations of DOX. We observed dose-dependent expression of both transgenes (m*Cherry* and *fLuc*) at 5 dpt (**Fig. 6A&B**). Notably, at 2 µM, DOX augmented rAAV2.5T transduction by ∼100-fold compared to the vehicle (DMSO) control (**Fig. 6B**).

**Fig. 6.**
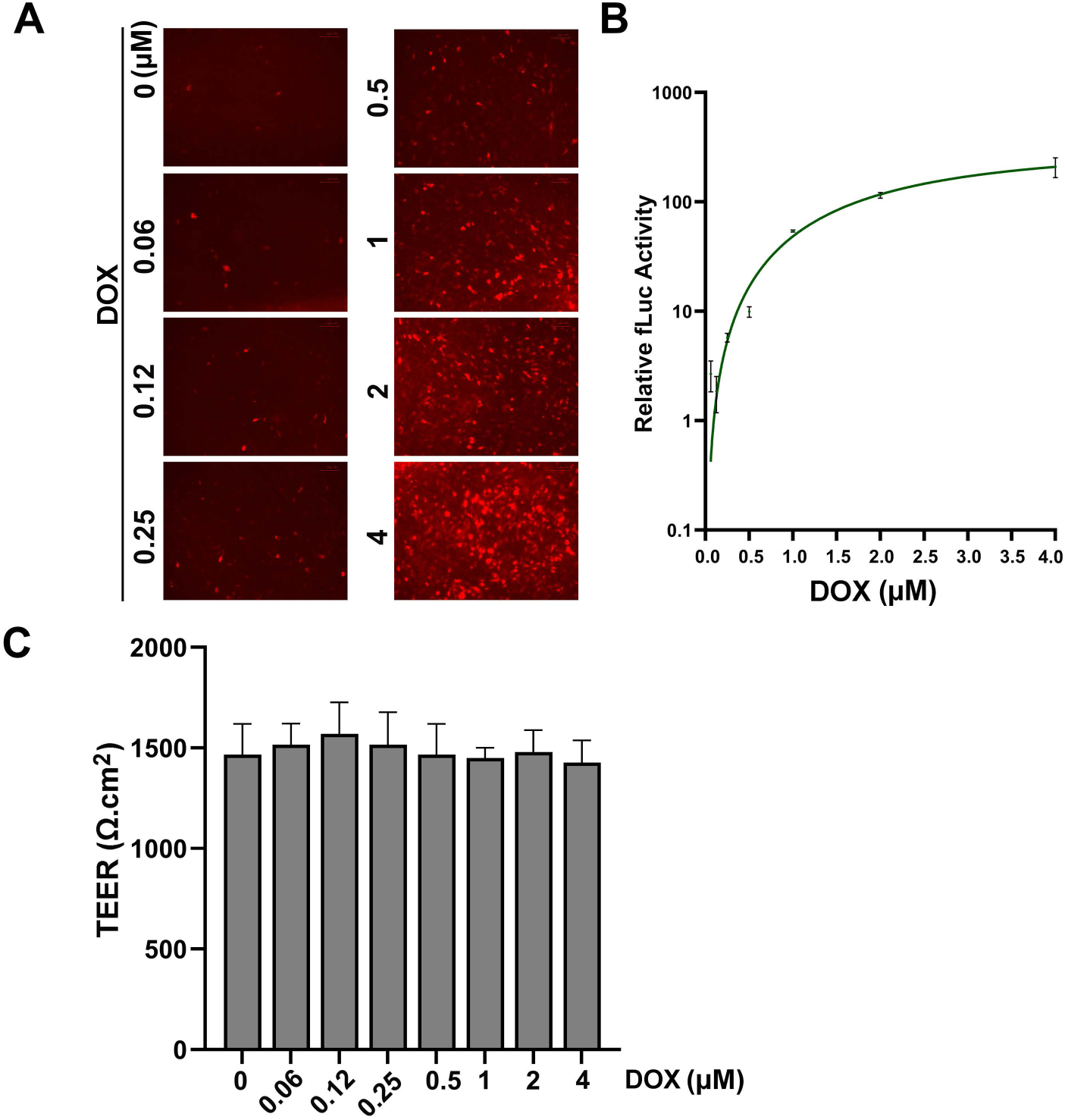
Doxorubicin enhances rAAV2.5T transduction in HAE-ALI cultures in a dose-dependent manner without disrupting epithelial barrier integrity. HAE-ALI cultures were treated with increasing concentrations of DOX as indicated for 16 h during apical transduction with rAAV2.5T at an MOI of 20,000 DRP/cell. **(A) mCherry expression.** Shown are representative fluorescence Images acquired at 5 dpt using an inverted fluorescence microscope (Nikon Ti-S). **(B) Quantification of luciferase activity.** fLuc activity was measured at 5 dpt and normalized to the DMSO-treated (0) control group, which was set to 1. **(C) Measurement of transepithelial electrical resistance (TEER).** TEER values were measured at 5 dpt to evaluate epithelial barrier integrity. Data are presented as mean ± SD. Statistical analysis was performed using one-way ANOVA followed by Dunnett’s multiple-comparison test. No statistically significant differences were observed between treated and untreated groups.

Importantly, DOX treatment did not compromise the epithelial barrier function of HAE-ALI cultures even at 4 µM, as evidenced by the absence of significant changes in transepithelial electrical resistance (TEER) (**Fig. 6C**).

Remarkably, DOX also completely compensated for the loss of GPR108 in rAAV2.5T transduction of HAE-ALI^GPR-KO^ cultures. Without DOX treatment, rAAV2.5T transduction yielded poor mCherry expression in HAE-ALI^NT^ and virtually undetectable expression in HAE-ALI^GPR-KO^ cultures [**Fig. 7A**, DOX(-)]. However, when transduction was performed in the presence of DOX, both cultures exhibited similarly high levels of mCherry expression [**Fig. 7A**, DOX(+)].

**Fig. 7.**
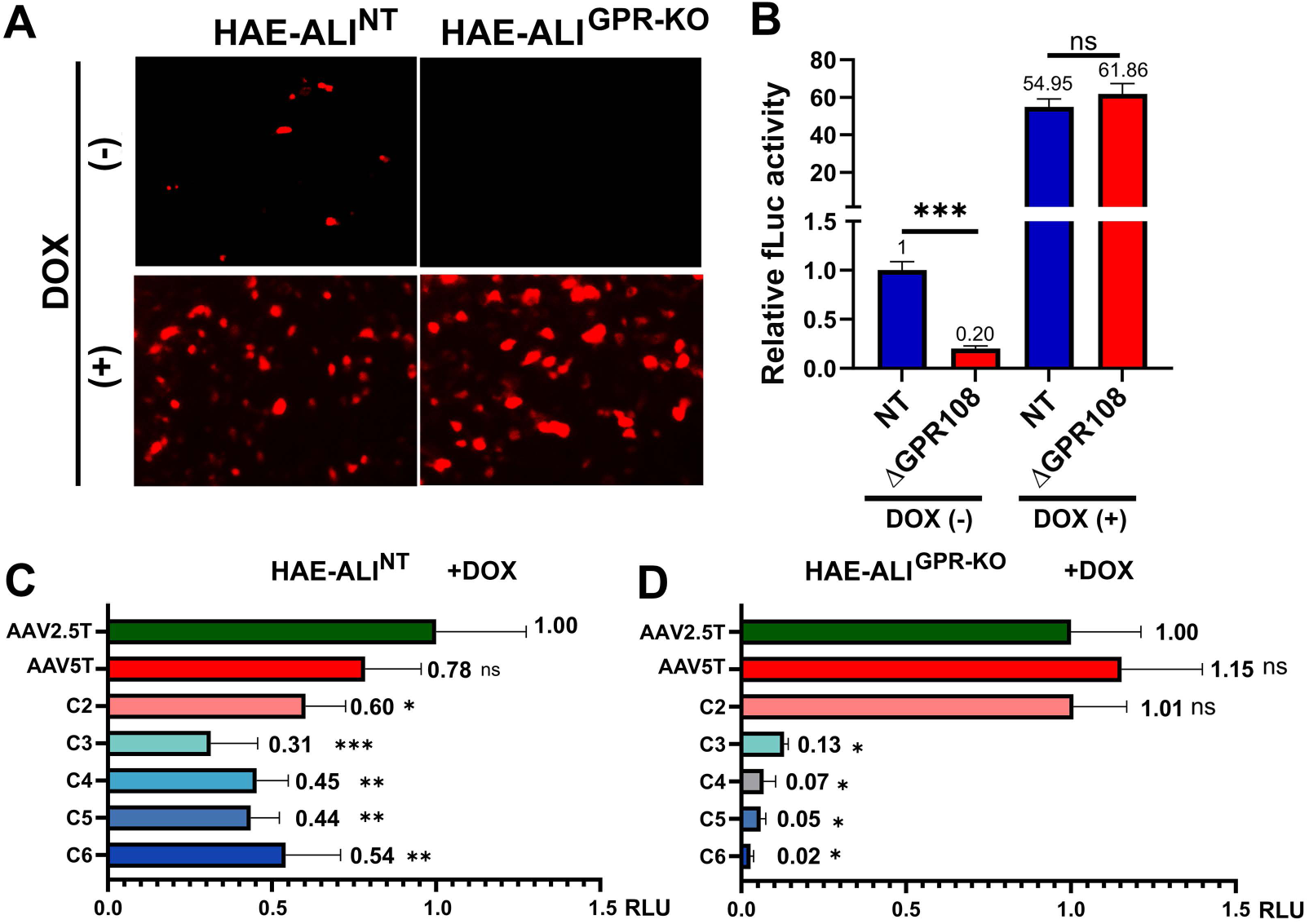
Doxorubicin compensates for the loss of GPR108 during rAAV2.5T transduction in HAE-ALI cultures. **(A&B) rAAV2.5T transduction in NT and GPR108-KO HAE-ALI cultures.** HAE-ALI cultures were transduced apically with rAAV2.5T at an MOI of 20,000 DRP/cell with or without 2 μM DOX. Transduction efficiency was assessed at 5 dpt. Representative mCherry fluorescence images were acquired using a Nikon Ti-S fluorescence microscope (A), and luciferase activity was quantified (B). fLuc activity was normalized to the untreated HAE-ALI^NT^ group, which was set to 1. Data are presented as mean ± SD from three independent experiments. Statistical analysis was performed using two-way ANOVA followed by Tukey’s multiple-comparison test. ***, P < 0.001; ns, not significant. **(C&D) Transduction of AAV2.5T, AAV5T, and VP1u chimeric mutants in polarized HAE-ALI cultures in the presence of DOX.** HAE-ALI^NT^ (C) and HAE-ALI^GPR-KO^ (D) cultures were transduced apically with the indicated rAAV vectors at an MOI of 20,000 DRP/cell in the presence of 2 µM DOX. At 5 dpt, luciferase activity was measured and was normalized to rAAV2.5T transduction, which was set to 1.0. Data are presented as mean ± SD from three independent replicates. Statistical significance was determined using one-way ANOVA followed by Dunnett’s multiple-comparison test. *, P < 0.05; **, P < 0.01; ***, P < 0.001; ns, no significant.

Quantitative analyses of fLuc activity mirrored these findings: without DOX, transduction in the GPR108-KO ALI cultures was reduced by 80% compared to the NT ALI cultures [**Fig. 7B**, DOX(-)]. In contrast, the addition of 2 µM DOX brought fLuc activity in both the NT control and GPR108-KO cultures to comparable levels, representing 55-fold and 310-fold increases relative to their respective untreated transduction [**Fig. 7B**, DOX(+)]. This finding is starkly different from that in AAVR-KO, where DOX treatment failed to exert a comparable compensatory effect on rAAV2.5T transduction in AAVR-KO ALI cultures (28). These data demonstrate that DOX treatment rescues the transduction deficit imposed by GPR108 deficiency and fully compensates for the GPR108 deficiency transduction in HAE-ALI cultures.

We further tested the DOX-responsiveness of AAV5T and its VP1u mutants (C2-C6) in HAE-ALI. In the presence of DOX, AAV5T and C2 showed 78% and 60%, respectively, of fLuc activity in transduction of HAE-ALI, compared with that of AAV2.5T; C3-C6 mutants also had 31% to 54% of fLuc activity in transduction, which correlated with the length of the substituted AAV2 VP1 sequences, compared with the AAV2.5T (**Fig. 7C**). Although DOX augmented transduction of AAV2.5T, AAV5T and C2 at the same level in HAE-ALI^GPR-KO^, surprisingly DOX failed to enhance transduction of C3-C6 mutants (**Fig. 7D**).

Taken together, these data demonstrated that DOX treatment fully compensated for the loss of GPR108 during transduction by rAAV2.5T, AAV5T, and the AAV5T.C2 mutant that has only 4 amino acid substitutions in VP1u, but not by the mutants containing 15 or more aa replacements from AAV2 VP1u. These findings strongly suggest that DOX-mediated enhancement requires a nearly intact VP1u in either AAV2 or AAV5.

### DOX-mediated rescue of rAAV2.5T transduction in GPR108-KO cells is independent of vector internalization and DNA damage response induction

We next investigated the mechanism underlying the DOX enhancement of rAAV2.5T transduction in HAE-ALI cultures. We hypothesized four possible pathways through which DOX enhances rAAV2.5T transduction: 1) facilitating the internalization of the vector into cells; 2) facilitating the import of vectors from the TGN to the nucleus; and 3) inhibiting proteasome activity, thereby reducing degradation of vectors in proteasomes; 4) inducing a DNA damage response (DDR) that in turn enhances rAAV2.5T transgene expression (**Fig. 8**).

**Fig. 8.**
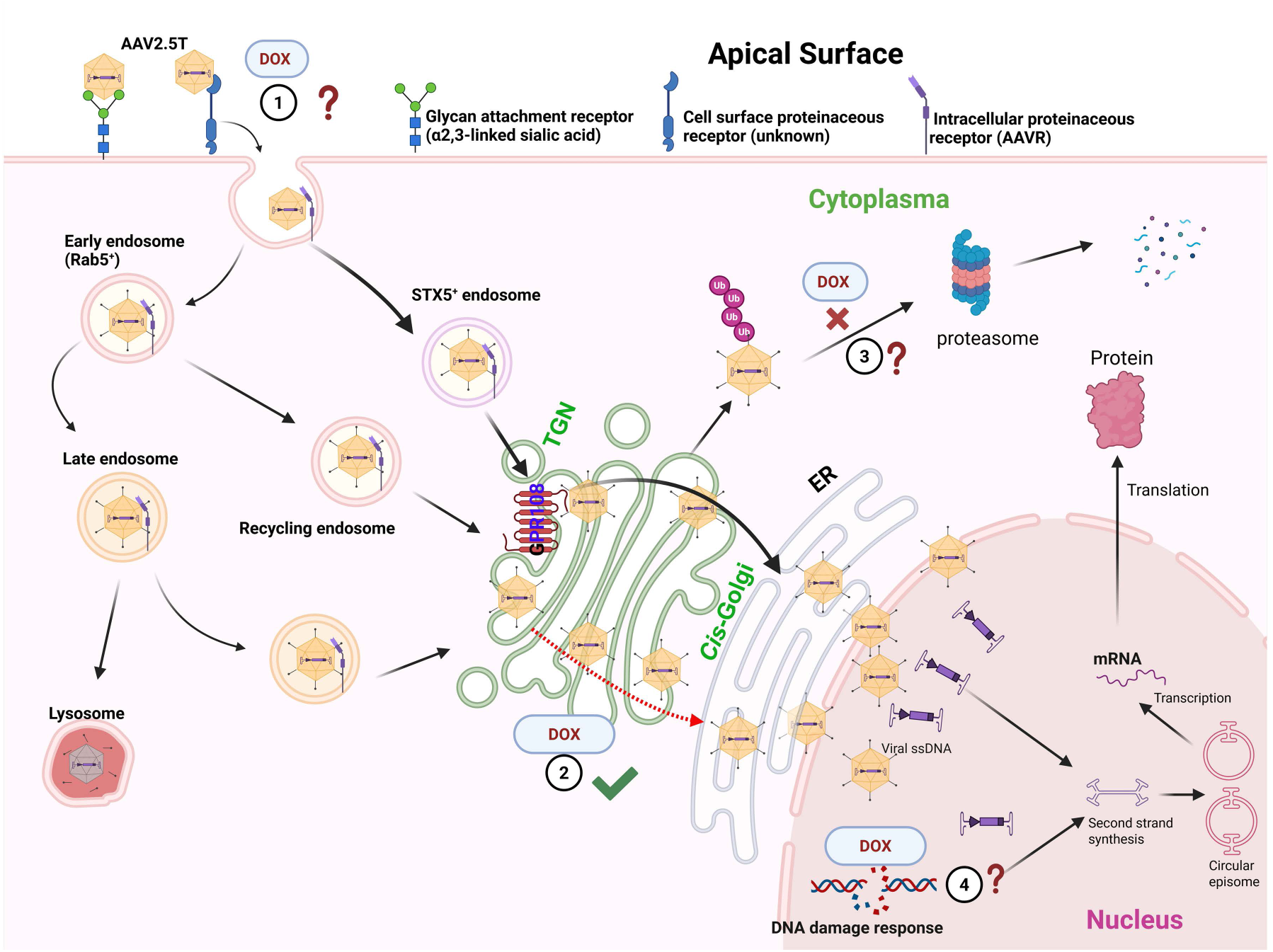
Proposed pathways by which doxorubicin increases rAAV2.5T transduction in HAE-ALI. Schematic illustration shows the proposed intracellular trafficking pathway of rAAV2.5T and potential mechanisms by which DOX enhances vector transduction. rAAV2.5T initially binds to cell-surface glycans containing α2,3-linked sialic acid and an unidentified proteinaceous cell surface receptor prior to endocytosis. Following internalization, vectors interact with AAVR, which is not required for vector internalization but required for intracellular trafficking, and traffic through multiple endosomal compartments, including STX5-positive retrograde endosomes and other endosomes. Acidification within endosomes promotes extrusion of VP1u from the capsid surface. A subset of vectors traffics to the *trans*-Golgi network (TGN) through the AAVR–TBC1D23-mediated retrograde transport pathway (17), where capsids interact with GPR108 within the TGN lumen to facilitate subsequent nuclear entry. DOX may enhance rAAV2.5T transduction through several non-mutually exclusive mechanisms: (1) increasing vector internalization or early endocytic trafficking; (2) bypassing or compensating for the GPR108-dependent trafficking step that promotes vector transport from the TGN toward the nucleus; (3) reducing proteasome-mediated capsid degradation; and/or (4) activating DDR pathways that facilitate second-strand synthesis, episome formation, or transgene expression after nuclear entry. After nuclear import, the viral ssDNA genome undergoes uncoating and second-strand synthesis to form transcriptionally active transduction intermediates, which are subsequently converted into episomal genomes, leading to persistent transgene expression. The schematic was created using BioRender.

First, we conducted a vector internalization assay with rAAV2.5T in NT and GPR108-KO ALI cultures, treated or untreated with DOX. We found no significant difference in vector internalization between the groups (**Fig. 9A**). Next, we investigated whether DOX increased rAAV2.5T transduction by inducing a DDR. We used DOX to treat HAE-ALI at concentrations of 1, 2, 4, and 10 µM, with HU (hydroxyurea) as a positive control. Western blotting detected gamma (γ)H2AX in the HU-treated culture and the 10 µM DOX-treated cultures, but not in the cultures treated with lower concentrations (1, 2, and 4 µM) (**Fig. 9B**). RPA2 phosphorylation was also not induced when the cultures were treated with lower DOX doses (**Fig. 9B**). Furthermore, we conducted a comet assay to detect damaged DNA in HAE-ALI treated with 2, 4, or 10 µM of DOX, and found no detectable evidence of DNA damage in cells of HAE-ALI treated with DOX at <4 µM, which was noted in the control cultures treated with H_2_O_2_ (**Fig. 9C**).

**Fig. 9.**
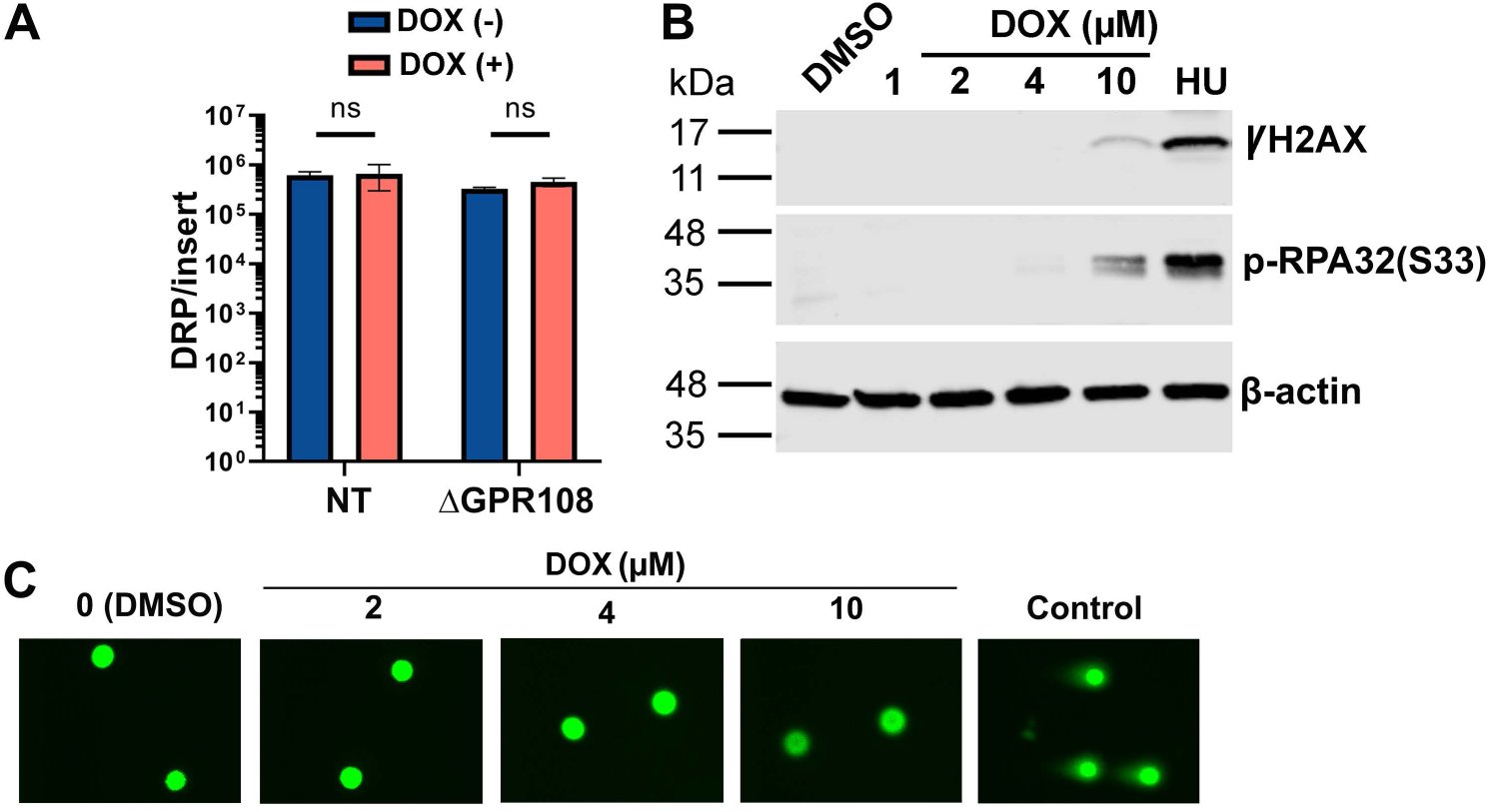
Doxorubicin-enhanced rAAV2.5T transduction is independent of vector internalization and DNA damage signaling. **(A) Vector internalization assay.** HAE-ALI cultures of NT or GPR108-KO were apically transduced with rAAV2.5T at an MOI of 20,000 DRP/cell in the presence or absence of 2 μM DOX. At 2 hpt, surface-bound vectors were removed and internalized vector genomes were quantified by qPCR. Data are presented as mean ± SD from three independent experiments. Statistical analysis was performed using two-way ANOVA followed by Tukey’s multiple-comparison test. ns, no significant difference. **(B) Analysis of DNA damage response markers following DOX treatment in HAE-ALI cultures.** Polarized HAE-ALI cultures were treated with increasing concentrations of DOX (1, 2, 4, or 10 μM) for 16 h. Hydroxyurea (HU, 2 mM) was included as a positive control to activate DDR. At 5 dpt, cells were harvested for Western blot analysis of γH2AX expression and phosphorylated RPA32 (p-RPA32), respectively. β-actin served as a loading control. **(C) Alkaline comet assay analysis of DNA damage following DOX treatment**. HAE-ALI cultures were treated with DMSO, 2, 4, or 10 μM DOX for 16 h. At 5 dpt, cells were subjected to alkaline comet assay analysis. H₂O₂ treatment (100 μM) served as the positive control. Representative images are shown.

Collectively, these results demonstrate that GPR108-KO does not impair AAV internalization and that low-dose DOX treatment does not significantly induce a DDR or damaged DNA in HAE-ALI cultures.

### DOX promotes AAV nuclear import independent of proteasome inhibition

It has been reported that DOX can inhibit proteasome activity (39) and that co-administration of DOX and proteasome inhibitor LLnL synergistically enhanced rAAV2 and rAAV5 transduction in HAE-ALI by preventing vector degradation in the cytosol (1), allowing more vectors to enter the nucleus, which results in high expression of the transgene delivered by rAAV vectors. Given that DOX is a pleiotropic compound, we investigated whether its enhancement of rAAV2.5T transduction is mediated through modulation of the ubiquitin-proteasome system. To address this, we measured proteasome activity in DOX-treated HAE-ALI cultures, with LLnL-treated cultures as a positive control. Treatment with LLnL at 40 µM for 16h significantly decreased proteasome activity by ∼90% but increased fLuc activity only by ∼2-fold (**Fig. 10A&B**), indicating that proteasome inhibition by LLnL only moderately increases rAAV transduction in HAE-ALI. In contrast, 2 µM DOX did not significantly reduce proteasome activity compared to the DMSO control, yet DOX treatment increased luciferase activity by 70-fold (**Fig. 10B**). Furthermore, combining DOX and LLnL did not significantly increase the transduction, compared with DOX alone (**Fig. 10B**). These data demonstrated that at a low concentration (2 µM) required for effective transduction, DOX plays a negligible role in inhibiting proteasome activity.

**Fig. 10.**
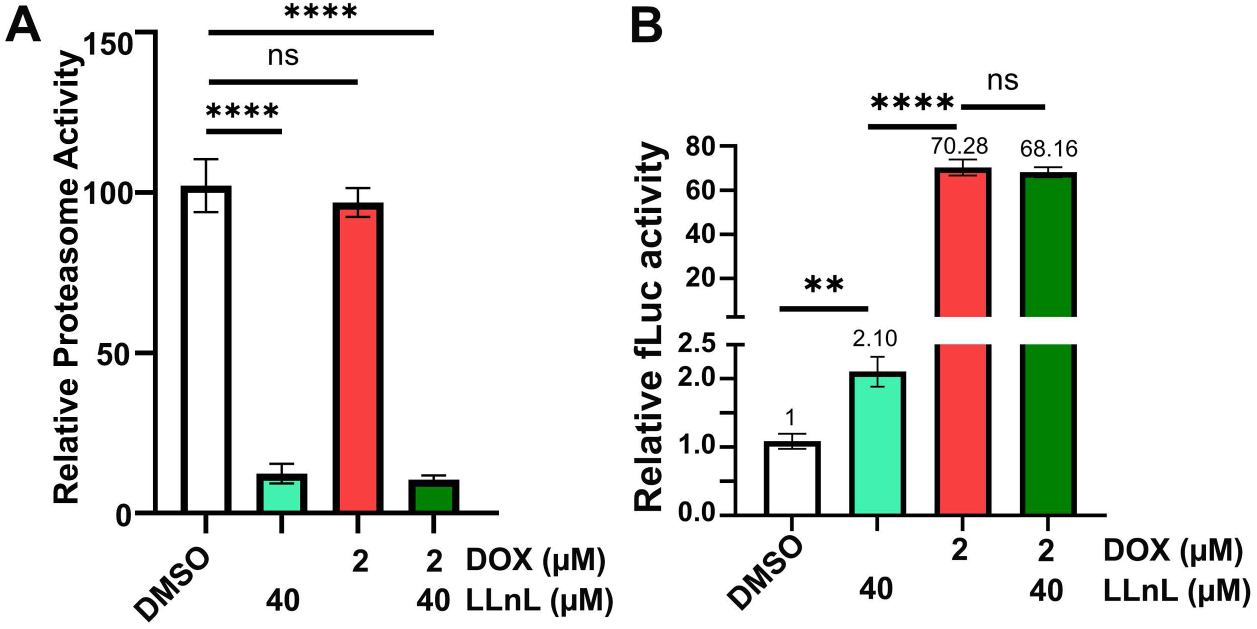
Doxorubicin enhances rAAV2.5T transduction independently of proteasome inhibition in polarized HAE-ALI cultures. **(A) Proteasome activity assay.** HAE-ALI cultures were treated with the proteasome inhibitor LLnL (40 μM), doxorubicin (DOX at 2 µM), or DOX (2 µM) plus LLnL (40 μM) for 16 h. DMSO-treated cultures served as controls. After treatment, cells were lysed, and proteasome activity was measured. **(B) Luciferase activity assay.** Polarized HAE-ALI cultures were apically transduced with rAAV2.5T at an MOI of 20,000 DRP/cell in the presence of LLnL (40 μM), Dox (2 μM), or combined LLnL (40 µM) and DOX (2 µM) treatment for ∼16 h. Luciferase activity was measured at 5 dpt and normalized to the DMSO-treated cultures. Data are shown as mean ± SD from three independent experiments. Statistical analysis was performed using one-way ANOVA followed by Tukey’s multiple-comparison test. **, P < 0.01; ****, P < 0.0001; ns, not significant.

Lastly, we investigated whether DOX functions by facilitating vector nuclear import. First, we performed immunofluorescent staining of AAV2.5T capsids in HAE-ALI^NT^ and HAE-ALI^GPR-KO^ cultures, and in HAE-ALI^AAVR-KO^ as a control. Notably, in the absence of DOX, some vectors (capsids) were detectable in the nuclei of NT cells [**Fig. 11A**, DOX(-)], whereas few to no capsids were observed in the nuclei of both GPR108-KO and AAVR-KO cells [**Fig. 11B-C**, DOX(-)]. Upon DOX treatment, however, nuclear capsid accumulation was markedly increased in cells of both GPR108-KO and NT ALI cultures but not of AAVR-KO ALI cultures [**Fig. 11B-C**, DOX(+)]. This finding suggests that DOX treatment facilitates AAV trafficking from the cytosol to the nucleus, effectively bypassing the requirement of GPR108. To corroborate this, we next conducted subcellular fractionation of rAAV2.5T-transduced HAE-ALI cultures (**Fig. 11D**) and quantified the nuclear and cytoplasmic viral DNA. Consistent with our imaging data, DOX treatment significantly increased the abundance of viral genome in the nuclei, with ∼4-fold increase in HAE-ALI^NT^, ∼25-fold increase in HAE-ALI^GPR-KO^ cultures, but had no effect on HAE-AL^AAVR-KO^ cells compared to their respective untreated controls (**Fig. 11E**).

**Fig. 11.**
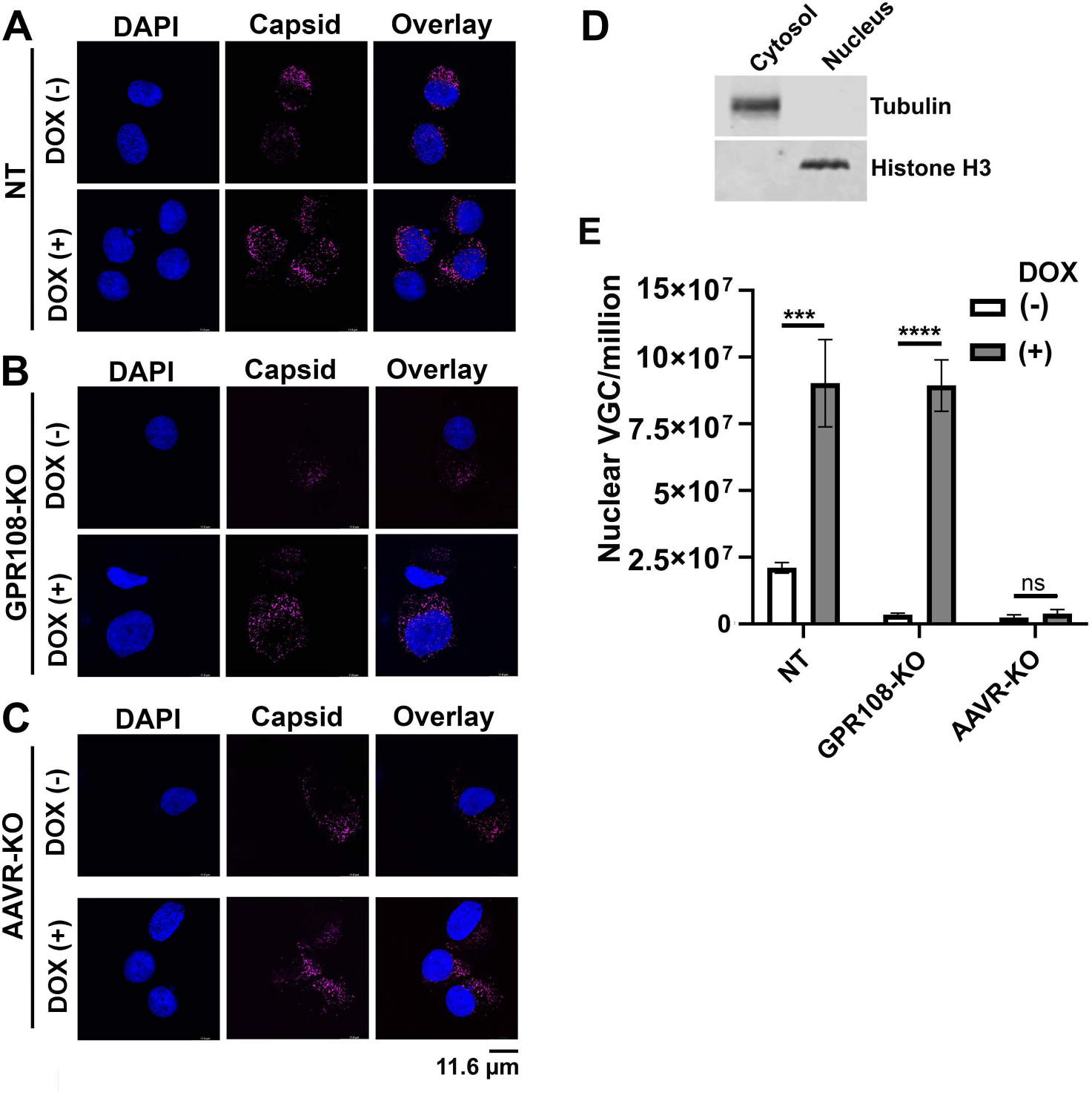
Doxorubicin promotes rAAV2.5T nuclear accumulation in HAE-ALI cultures in a GPR108-independent but AAVR-dependent manner. **(A-C) Immunofluorescence analysis of intracellular rAAV2.5T capsids.** HAE-ALI^NT^ (A), HAE-ALI^GPR-KO^ (B), and HAE-ALI^AAVR-KO^ (C) cultures were transduced apically with rAAV2.5T at an MOI of 20,000 DRP/cell in the absence or presence of doxorubicin (DOX, 2 μM). At 5 dpt, cells were dissociated from Transwell inserts, cytospun onto glass slides, and immunostained with anti-AAV5 capsid. Nuclei were counterstained with DAPI. Images were acquired using a Leica TCS SP8 STED confocal microscope with a 100× objective. Representative images are shown. Scale bar = 11.6 μm. **(D) Validation of nuclear and cytosolic fractionation.** Cytosolic and nuclear fractions isolated from HAE-ALI cultures were analyzed by Western blotting with antibodies against tubulin (cytosolic marker) and histone H3 (nuclear marker) to assess fraction purity. **(E) Quantification of nuclear viral genomes.** HAE-ALI^NT^ (A), HAE-ALI^GPR-KO^ (B), and HAE-ALI^AAVR-KO^ cultures were transduced with rAAV2.5T in the absence or presence of DOX (2 μM). At 16 hpt, cells were harvested and separated into nuclear and cytosolic fractions. DNA extracted from purified nuclei was subjected to qPCR quantification of vector genomes. Data are presented as mean ± SD from four independent experiments. Statistical significance was determined using an unpaired two-tailed Student’s t-test comparing control and DOX-treated groups within each genotype. ***, P < 0.001; ****, P < 0.0001; ns, not significant.

Taken together, these data indicated that DOX enhances transduction by promoting vector nuclear entry through a mechanism that bypasses GPR108 but remains dependent on AAVR.

## Discussion

rAAV2.5T exhibits higher tropism to polarized human airway epithelia than its parent AAV5 (27). Here, we revealed that rAAV2.5T transduction depends on GPR108, which is mainly determined by the N-terminal 15 aa of the VP1u in the AAV2.5T capsid. We also revealed that in polarized HAE cultures, DOX enhances rAAV2.5T transduction through an increase in vector nuclear entry, but not in vector internalization, inhibition of proteasome activity, or the induction of DDR.

### rAAV2.5T internalization and intracellular trafficking in HAE-ALI

AAV enters cells via receptor-mediated endocytosis, interacting with primary glycan-attachment receptors and proteinaceous co-receptors (21,42). AAV2.5T uses cell surface α2,3 N-linked sialic acid as the primary attachment receptor (43), but the proteinaceous receptor has not been identified (29) (**Fig. 8**). Once the vector is internalized, it traffics through different endosomal compartments, where the VP1u is extruded from the capsid surface (19,44). VP1u comprises the 137-aa N-terminal region unique to the large VP1. VP1u contains a PLA_2_ activity domain in the center (45) (**Fig. 4A**), which is essential for the establishment of a productive AAV infection (18,46).

Although VP1u is initially concealed within the capsid, its emergence on the virion surface facilitates penetration of the AAV capsid through the last membrane vesicle, e.g., the Golgi body or endoplasmic reticulum (ER), allowing its entry to the nucleus, where it uncoats and loads its genome to initiate replication/gene expression (**Fig. 8**). For most AAV serotypes, during internalization and trafficking to TGN, AAVR plays an important role (14,15). However, we found that AAVR is not necessary for the internalization of rAAV2.5T into polarized HAE through apical surface; instead, it plays a crucial role in its intracellular trafficking (28). Since AAVR is a type-I transmembrane protein cycling between the cell surface and the TGN (14), it functions in trafficking the vector from the endosomes to the TGN (17).

GPR108 is a poorly characterized seven-transmembrane protein that localizes to the Golgi apparatus, and its physiological function remains incompletely understood. GPR108 has been identified as a conserved essential factor for transduction of rAAV2 but not of rAAV5 (25,26).

Our study found that GPR108 is broadly distributed throughout the cytoplasm and colocalizes not only with plasmid-expressed VP1u domains of AAV2 and AAV5 but also with intact AAV2 and AAV5 capsids following vector transduction. TGN, defined by TGN46 staining, occupied a small perinuclear region nested within the broader GPR108-localized compartments. These observations suggest that GPR108 participates in AAV trafficking after capsids reach and enter the TGN. Since both AAV5 VP1u and AAV5 capsids colocalize robustly with GPR108, we speculate that a GPR108-counterpart may facilitate the downstream trafficking of AAV5 capsids from the TGN to the nucleus. Although a previous genome-wide screen using rAAV5 did not identify a functional counterpart to GPR108 (28), the possibility of its existence cannot be excluded and warrants further investigation.

### The N-terminus of AAV2 VP1u determines the GPR108 dependence of rAAV2.5T transduction

Our VP1u-engineered AAV5T mutants (C2-C6) helped us to better understand the function of the AAV2 VP1u in AAV2.5T transduction. From AAV5T.C3-C6 mutants, where the AAV2 VP1u is extended from 15-44 aa, we observed a corresponding increase in the transduction efficiency. AAV5T.C6 that contains 1-44 aa of AAV2 VP1u showed ∼80% transduction efficiency, compared to rAAV2.5T in HeLa cells and HAE-ALI. The AAV5T.C3 mutant significantly lost transduction in GPR108-KO cells, compared to the parent AAV2.5T and C2 mutant, suggesting that aa1-15 of the AAV2 VP1u is critical for the GPR108 dependence.

We also constructed a C7 variant that includes aa1-80 of AAV2 VP2 to achieve higher transduction than C6; however, the C7 mutant showed no transduction ability (**Fig. S3**). This indicates that the integrity of the VP1u PLA_2_ domain is important in rAAV transduction. The structures of the C2-C6 VP1u variants predicted by AlphaFold showed that the 4 α-helices are retained, suggesting the importance of the VP1u protein structure in rAAV transduction, which likely maintains the structure of PLA_2_, as well as the VP1u N-terminus, for their function.

Unfortunately, no VP1u structures of any parvovirus have been resolved to date (18). Notably, the VP1 N-terminal domain controls rAAV genome transcription through chromatin accessibility and histone modifications on the genomes (47). Non-mammalian AAVs cannot transduce human cells, but that transduction can be rescued with a 40-aa N-terminal domain from primate AAVs (47,48). Thus, the VP1u N-terminus functions not only in GPR108-dependent transduction, but also in the epitranscriptomic regulation of transgene expression.

### Mechanisms of DOX in the enhancement of rAAV2.5T transduction in HAE-ALI

AAV transduction can be significantly enhanced by DOX, particularly in HAE-ALI cultures. We ruled out the possibility that DOX enhances AAV internalization into cells. In addition, treatment of HAE-ALI cultures with 2 µM DOX did not induce detectable expression of γH2AX or phosphorylated RPA32 (49,50), indicating the absence of a significant DDR under these conditions. Our comet assay further supported that treatment with 2 µM DOX did not cause detectable damage to cellular DNA. We previously observed that AAV2.5T transduction in HAE-ALI cultures can induce a DDR (33); however, inhibition of the DDR paradoxically increases rAAV transduction. Therefore, any DDR potentially induced by low concentrations of DOX is unlikely to contribute to the DOX-mediated enhancement of rAAV transduction.

It has been proposed that DOX inhibits proteasome activity, thereby reducing vector disassembly and degradation in the cytosol. Consequently, more AAVs are transported into the nucleus and express the transgene (51) (**Fig. 8**). At low concentrations, e.g., 2 µM, DOX does not inhibit proteasome activity. This is consistent with a previous report that DOX affects cellular proteasome activity at higher concentrations (≥6.25 µM) (39). In addition, DOX treatment at 2 or 4 µM did not alter the GPR108 expression (**Fig. S2B**), suggesting that at these working concentrations, DOX is unlikely to alter host gene expression.

Lastly, we confirm that DOX treatment can significantly facilitate vector nuclear import. While GPR108-KO severely impairs rAAV2.5T transduction in HAE-ALI, transient DOX treatment at a low concentration compensates for the loss of rAAV transduction in GPR108-KO cells, but not in AAVR-KO cells, even though AAV2.5T successfully enters AAVR-KO HAE-ALI cultures (28). Given that AAVR is a key host factor required for trafficking AAV to the TGN (17), we hypothesize that DOX promotes vector trafficking from the TGN to the nucleus through a GPR108-independent pathway (**Fig. 8**, dashed arrow). Furthermore, we observed that DOX failed to enhance transduction of AAV5T.C3–C6 mutants containing chimeric VP1u regions, suggesting that this DOX-mediated enhancement depends on intact VP1u derived from either AAV2 or AAV5, rather than a chimeric construct. These findings warrant further investigation into the precise intracellular trafficking pathway that mediates vector transport from the TGN to the nucleus, with important implications for improving the clinical efficacy of rAAV-based gene therapy.

## Materials and Methods

### Cells and cell cultures

#### Cells

HeLa cells (ATCC #CCL-2) were grown in Dulbecco’s Modified Eagle Medium (DMEM; HyClone #SH30022.01, Cytiva, New York, NY) supplemented with fetal bovine serum (FBS) at 10% and penicillin-streptomycin at 100 units/mL in a humidified incubator with 5% CO_2_ at 37°C. CuFi-8 cells were immortalized from human primary airway epithelial cells isolated from a cystic fibrosis patient by expressing hTERT and HPV E6/E7 genes (52). They were cultured on collagen-coated 100-mm dishes in PneumaCult Ex Plus medium (#05040; StemCell, Vancouver, BC).

### Human airway epithelium (HAE) cultured at an air-liquid interface (ALI)

Proliferating CuFi-8 cells dissociated from flasks were directly loaded on collagen-coated Transwell permeable supports (#3460; Costar, Corning) with PneumaCult Ex Plus in both apical and basal chambers. One day after seeding, media were replaced with PneumaCult-ALI medium (#05001; StemCell) for 1–2 days, and then the media in the apical chamber were removed. The cells were polarized in PneumaCult-ALI medium at an ALI for 3–4 weeks (53).

The maturation of the polarized HAE-ALI cultures was determined by the transepithelial electrical resistance (TEER) measured with a Millicell ERS-2 volt-ohm meter (MilliporeSigma, Burlington, MA). HAE-ALI cultures with a TEER value of >1,000 Ω·cm^2^ were used for experiments.

### Production of rAAV vectors

The dual-reporter rAAV vectors (rAAV.fLuc-mCherry) were produced using a triple-plasmid transfection method using PEImax (Cat# 24765-2, MW 40,000, Polysciences, Inc.) (54). The AAV trans-helper plasmids include AAV2Rep-AAV2 capsid expressing plasmid, AAV2Rep-AAV5 capsid expression plasmid, AAV2 Rep and AAV2.5T capsid-expressing plasmid pAV2.5Trepcap (27,34,35), and the AAV2 Rep and chimeric (mutant) AAV5T capsid-expressing plasmids (as described below). In brief, an AAV trans-helper plasmid, the adenovirus 2 helper pHelper, and the AAV2 transgene plasmid pAV2F5tg83luc-CMVmCherry (55) were transfected at a molar ratio of 1:1:1 into HEK293 cells cultured in 150-mm plates. rAAV vectors were purified by density gradient ultracentrifugation in cesium chloride, following a method described previously (51). Vector titers were determined by real-time quantitative polymerase chain reaction (qPCR) using primers and probes specific to the *mCherry* transgene as DNase I-resistant particle (DRP) per ml (55).

### Plasmid constructions

#### Flag-tagged VP1u expression plasmids

pCMV-AAV2VP1u^Flag^ and pCMV-AAV5VP1u^Flag^ were constructed by cloning the unique region of the large capsid protein (VP1u)-coding sequences of AAV2 and AAV5, tagged with Flag at the C-terminus, into pcDNA3 through HindIII and EcoRV sites.

#### Chimeric AAV5T capsid-expressing plasmids

pAV2Rep-AV5T was constructed by changing the VP1u to be AAV5 in pAV2.5Trepcap. Then, based on pAV2Rep-AV5T, the AAV5 VP1u sequence was mutated to AAV2 aa13-16 (C2), aa1-15 (C3), aa1-23 (C4), aa1-32 (C5), and aa1-44 (C6) to construct pAV2Rep-AV5T.C2-C6, respectively.

### Plasmid transfection

HeLa cells were transfected with pCMV-AAV2VP1u^Flag^ and pCMV-AAV5VP1u^Flag^, respectively, using the PEImax transfection reagent at a DNA: PEI ratio of 1:3 in Opti-MEM (Invitrogen). The total amounts of plasmid DNA were kept constant in each group by supplementation with an empty vector.

### Vector internalization assay

HAE-ALI cultures were infected with rAAV2.5T at an MOI of 20,000 DRP/cell from the apical side. HeLa cells were infected at the same MOI. The cultures were then incubated in a humidified incubator with 5% CO_2_ at 37°C for 2 hours, followed by three D-PBS washes to remove unbound vectors. We carried out vector internalization assay as previously described (56). In brief, the cultures were washed with Accutase (#AT104, Innovative Cell Technologies, Inc.) three times and then incubated with Accutase at 37°C for 1 hour. After washing with PBS three times, viral DNA was collected using Quick-DNA/RNA Pathogen Kits (Zymo Research, #R1042) and quantified by qPCR using *mCherry* primers/probe.

### Proteasome activity assay

The Proteasome Activity Assay Kit (Fluorometric) (Abcam, #ab107921) was used to determine proteasome activities in cells. Briefly, 2 x 10^6^ cells were harvested and washed with cold PBS three times. The cell pellets were resuspended in 1 mL of 0.5% NP-40 in dH_2_O and were quickly homogenized by pipetting up and down a few times. Then, the cell lysates were centrifuged for 10–15 minutes at 4°C at 13,000 g, and the supernatant was transferred to a clean tube. Four types of samples were prepared as follows: Standard wells = 100 μL of standard dilutions; Sample wells = 50 μL of samples in duplicate (adjust the volume to 100 μL/well with Assay Buffer); Positive Control wells = 10 μL of Positive Control III, and Background wells = 100 μL of Assay Buffer. To all samples and positive control wells, 1 μL of Proteasome substrate was added, and they were protected from light. The mixture was thoroughly mixed and then incubated at 37°C, still protected from light. Subsequently, fluorescence was measured on a microplate reader (BioTek Synergy H1) at the initial time point (T1; Ex/Em = 350/440 nm). The reactions were then incubated at 37°C for 30 min in the dark, after which fluorescence was measured again at the final time point (T2; Ex/Em = 350/440 nm).

Proteasome activity was calculated from the increase in fluorescence between T1 and T2 according to the manufacturer’s instructions.

### Comet assay

The comet assay was performed using a kit purchased from Cell Biolabs Inc. (#STA-351, San Diego, CA) according to the manufacturer’s instructions as previously described (57). In brief, cells in the Transwells of HAE-ALI cultures treated with DOX at 2, 4, or 10 µM in the basolateral chamber for 16 hours or with 100 mM H_2_O_2_ were used as a positive. Cells were dissociated from the supportive membrane with trypsin and diluted in PBS. The cells were mixed with 1% low-melting-point agarose and transferred onto slides. After fixing at 4°C, the slides were electrophoresed in alkaline buffer and stained with Vista green dye. The images were captured under a Nikon Eclipse C1 Plus inverted microscope.

### Cell fractionation

The NE-PER Nuclear and Cytoplasmic Extraction Reagents (ThermoFisher, #78833) were used to fractionate cytosol and nuclei. Briefly, cells were dissociated from the flask/transwell, centrifuged at 500 × g for 5 minutes, and washed with cold PBS three times. Then, the cell pellet was centrifuged at 500 × g for 2-3 minutes. The supernatant was removed carefully, and the cell pellet was kept as dry as possible. Next, 200 μL of ice-cold CER I (Cytoplasmic Extraction Reagent I) was added to the cell pellet, and the tube was vigorously vortexed on the highest setting for 15 s to fully suspend the cell pellet. The tube was then incubated on ice for 10 minutes. Afterward, 11 μL of ice-cold CER II (Cytoplasmic Extraction Reagent II) was added to the tube, and the tube was vortexed for 5 seconds on the highest setting before being set on ice for 1 minute. All the tubes were centrifuged for 5 minutes at maximum speed in a microcentrifuge (∼16,000 × g). The supernatant (cytoplasmic extract) was immediately transferred to a clean, pre-chilled tube. The insoluble pellet contained the nucleus fraction. To each tube, 100 μL of ice-cold NER (Nuclear Extraction Reagent) was added, and they were vortexed at the highest setting for 15 seconds. Then, all the tubes were placed on ice and vortexed for 15 seconds every 10 minutes, for a total of 40 minutes. The tubes were centrifuged at maximum speed (∼16,000 × g) for 10 minutes. Subsequently, the supernatant (nuclear extract) fractions were immediately transferred to clean, pre-chilled tubes.

### Sodium dodecyl sulfate–polyacrylamide gel electrophoresis (SDS-PAGE) and Western blotting

Cells were dissociated from the flask/transwell and solubilized with RIPA buffer [150 mM NaCl, 1% NP-40, 0.5% sodium deoxycholate, 0.1% SDS, 50 mM Tris-HCl, pH 7.4]. The cell lysates were mixed with 5 × Laemmli protein loading buffer and boiled for 10 min. Then, the samples were subjected to SDS-10% PAGE, followed by transferring the separated proteins onto a nitrocellulose membrane. After blocking with non-fat milk for 1 hour, the membrane was incubated with the primary antibody overnight, followed by incubation with an infrared dye-conjugated IgG (H+L) secondary antibody for 1 hour. Finally, the membrane was imaged on a Li-Cor Odyssey imager (LI-COR Biosciences, Lincoln, NB).

### rAAV transduction

For transduction of HeLa cells, cells were seeded overnight in 48-well plates. rAAV was added into each well at an MOI of 20,000 DRP/cell. The transduction efficiency was analyzed using a firefly luciferase assay and mCherry intensity at 3 days post-transduction.

For apical transduction of HAE-ALI, 400 µL of PBS-diluted rAAV vector was added into the apical chamber of the Transwell (Coster #3460) at an MOI of 20,000 DRP/cell. Then, 1 mL of culture media in the presence or absence of DOX at 2 µM, unless indicated, was added to the basolateral chamber. After 16 hours, all liquid in the apical and basolateral chambers was removed and washed with D-PBS, pH7.4 (Corning) three times. Fresh culture media (without DOX) were then added to the basolateral chamber.

### Luciferase assays

Firefly luciferase (fLuc) activity was detected using the Luciferase Assay System (Promega, #E4550). HeLa cultured at 48-well plates or HAE-ALI were transduced with rAAV. At 3 dpt, cell culture media were removed, and the cells were lysed in 200 µL of Lysis Buffer (Promega), then frozen at -80°C. After thawing, firefly luciferase expression was measured in relative light units (RLU) on a multi-mode microplate reader using (Synergy H1, BioTek).

### Generation of gene knockout stable cell lines

#### pLentiCRISPRv2 constructs

sgRNAs targeting GPR108 (5’-GTG GGG GCA GCG GCT ACT TC-3’), were cloned in plentiCRISPRv2 vector (54). The lentiCRISPRv2 expressing a scramble sgRNA (5’-GTA TTA CTG ATA TTG GTG GG-3’) was used as a non-target (NT) control, sgRNAs targeting AAVR was described in a previous publication (28).

#### gRNA-based lentivirus production

Lentivirus was produced in HEK293FT cells by transient transfection using PEI Max and transduced to cells according to a previously published protocol (54).

#### Generation of GPR108 knockout cell lines

HeLa or CuFi-8 cells were seeded at a density of 1 × 10^5^ cells per well in a 24-well plate. For CuFi-8 cells, the plates were collagen-coated. When the cells reached 80% confluence, the lentivirus containing GPR108 sgRNA was added to each well at ∼5 MOI. At two days post-transduction, puromycin was added at 2 µg/ml to select for gene knockout cells. The selected cells were then digested and seeded into a 96-well plate at a density of 0.5 cell/well. Once sufficient confluence was reached, the cells in each well were reseeded into a 6-well plate, and Western blot analysis was performed to determine the knockout efficiency.

### Immunofluorescence confocal microscopy

For rAAV-transduced cells of HAE-ALI cultures, the cells were dissociated off the supportive membranes of the Transwell inserts by incubation with Accutase. After incubation for 1 hour at 37°C, cells were completely detached from the membrane and well separated. The cells were then cytocentrifuged onto slides at 1,500 rpm for 3 minutes in a Shandon Cytospin 3 cytocentrifuge. The slides were fixed with 4% paraformaldehyde in D-PBS for 10 minutes and washed with D-PBS for 5 minutes three times. The fixed cells were permeabilized with 0.5% Triton X-100 for 15 min at room temperature. Then, the slide was incubated with one or two primary antibodies in D-PBS with 2% FBS for 1 h at 37°C. After washing three times, the slide was incubated with Alexa 488 (Green) and/or Alexa 594 (Red)-conjugated secondary antibodies, followed by staining of the nuclei with DAPI.

The stained slides were visualized under a Leica TCS SP8 or Nikon CSU-W1 SoRa confocal microscope at the Confocal Core Facility of the University of Kansas Medical Center. Images were processed with the Leica Application Suite X software.

### Phospholipase A2 (PLA_2_) activity assay

rAAV vectors (1 x 10^11^ DRP) were subjected to 80°C heating for 15 minutes to externalize VP1u of AAV5T-based capsids (58), and then assayed for PLA_2_ enzymatic activity using the Secretory Phospholipase A2 Assay Kit (Cayman, Ann Arbor, MI) (59,60). The assay was carried out following the manufacturer’s recommended protocol but with extended incubation periods (∼3 h) before absorbance (405 nm) readings (61).

### Antibodies used in the study

The following antibodies were used in this study: anti-Flag (F2555; Sigma), TGN46 (#NBP1-19643SS; Novus), GPR108 (#A16554; Abclonal), anti-tubulin (#A12289, Abclonal), anti-Histone H3 (#A2348, Abclonal), anti-AAV5 capsid (#03-615148; ARP), anti-AAV2 capsid (#61055; ARP), phospho-H2AX-S139 (γH2AX; #05-636; Millipore-Sigma), phopsho-RPA2-Ser33 (#AP1479; Abclonal), and β-actin (clone AC-15; Millipore-Sigma).

### Statistical analysis

All data are presented as mean ± standard deviation (SD) from at least three independent experiments and were analyzed using GraphPad Prism software. Error bars represent mean ± SD. Statistical significance between two groups was determined using an unpaired two-tailed Student’s t-test. For comparisons involving multiple groups, one-way or two-way analysis of variance (ANOVA) followed by the appropriate multiple-comparison test, as specified in the figure legends, was performed. P values < 0.05 were considered statistically significant. Statistical significance is indicated as follows: ****, P < 0.0001; ***, P < 0.001; **, P < 0.01; *, P < 0.05; and ns, not significant.

## Data availability

All data used to evaluate the conclusions in this study are presented in the paper and the **Supplementary Materials**.

## Acknowledgments

The study was supported by NIH grants R01HL174593 (J.Q. and Z.Y.), AI182645 (J.Q. and Z.Y.) and AI180416 (J.Q.), as well as Cystic Fibrosis Foundation YAN23G0 (Z.Y.). We acknowledge the Imaging Core Laboratory at the University of Kansas Medical Center for the use of the super resolution confocal microscope Leica SP8 STED, supported by NIH S10 OD 023625, and the Nikon CSU-W1 SoRa, supported by NIH S10 OD 032207. The funders had no role in study design, data collection and interpretation, or the decision to submit the work for publication.

## Notes

### Competing Interest Statement

The authors have declared no competing interest.

