## Supplemental Figures and Legends for "AAV VP1 unique region (VP1u) determines GPR108 dependence for AAV transduction of human airway epithelium and its rescue by Doxorubicin"

### Supplemental Materials

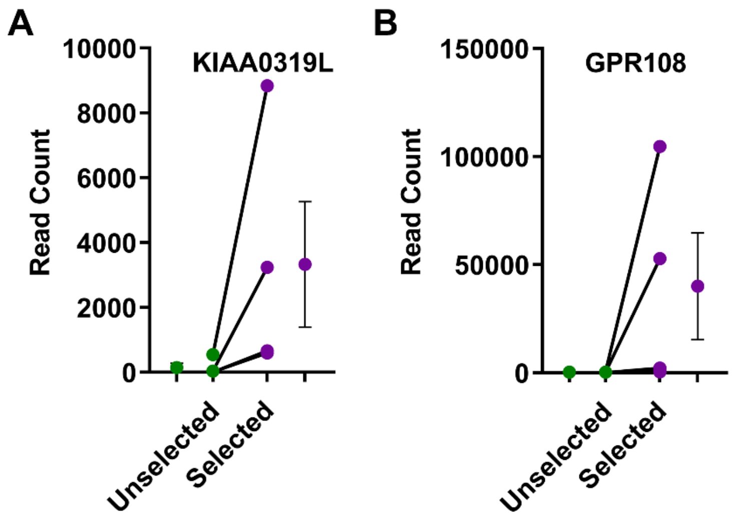


**Fig. S1. Enrichment of AAVR (KIAA0319L) and GPR108 gRNAs in a genome-wide CRISPR knockout screen following rAAV2.5T transduction.**

A genome-wide CRISPR KO screen in HeLa S3 cells infected with rAAV2.5T carrying an mCherry reporter was performed previously (1). Cells transduced with the CRISPR gRNA library were subsequently infected with rAAV2.5T carrying an mCherry reporter, and mCherry-negative cells were isolated by fluorescence-activated cell sorting (FACS) for next-generation sequencing analysis of enriched gRNAs. Read counts of individual gRNAs targeting AAVR (**A**) and GPR108 (**B**) in unselected and selected cell populations are shown. Each dot represents one of four independent gRNAs targeting the indicated gene, and lines connect the corresponding gRNAs between unselected and selected populations. Data are presented as read counts from next-generation sequencing analysis. Error bars indicate mean ± SD of the four gRNAs targeting each gene.

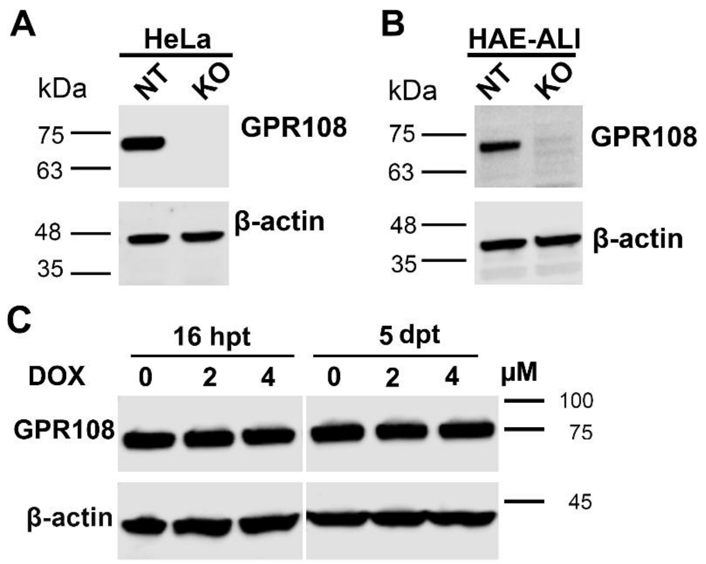


**Fig. S2. Validation of GPR108 knockout and the effect of doxorubicin on GPR108 protein expression.**

**(A&B) Validation of GPR108 knockout (KO).** The lysates of HeLa cells (A) and polarized HAE-ALI cultures (B) from non-targeting control (NT) and **GPR108**-knockout (KO) cells were analyzed by SDS-PAGE followed by immunoblotting with an anti-GPR108 antibody. β-actin served as the loading control. **(C) Doxorubicin (DOX) does not alter GPR108 protein expression in HAE-ALI cultures.** Cultures were treated with medium containing DMSO (0; no DOX vehicle control), 2, or 4 μM DOX in the basolateral compartment. At 16 h post-treatment (hpt), whole-cell lysates were collected. Remaining cultures were refreshed with basolateral medium without DOX and harvested as whole cell lysates at 5 days post-treatment (dpt). Cell lysates were analyzed by immunoblotting with an anti-GPR108 antibody. β-actin served as the loading control. Representative blots are shown. Molecular weight markers in kDa are indicated.

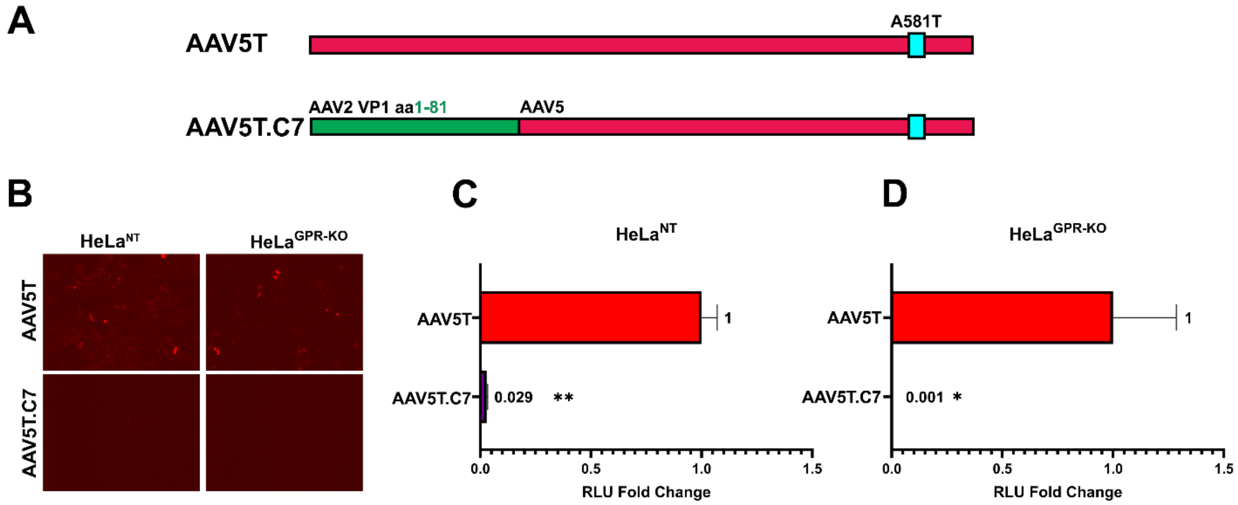


**Fig. S3. Replacement of the AAV2 VP1u N-terminal region (aa 1–81) severely impairs AAV5T transduction.**

**(A) Schematic of AAV5T and the C7 chimeric capsid**. In AAV5T.C7, amino acids 1–81 of the AAV5 VP1 unique region (VP1u) were replaced with the corresponding AAV2 sequence. AAV2-derived sequences are shown in green, AAV5-derived sequences in magenta, and the A581T mutation is indicated in cyan. **(B) mCherry expression.** Representative fluorescence images of HeLa$^{NT}$ and HeLa$^{GPR108\text{-}KO}$ cells transduced with AAV5T or the C7 chimera at an MOI of 20,000 DNase-resistant particles (DRP)/cell. Images were acquired at 5 days post-transduction (dpt). **(C&D) Quantification of firefly luciferase (fLuc) activity in HeLa$^{NT}$ (C) and HeLa$^{GPR108\text{-}KO}$ (D) cells**. Cells were transduced at an MOI of 20,000 DRP/cell, and fLuc activity was measured at 5 dpt and normalized to AAV5T. Data are presented as mean ± SD. Statistical significance was determined using an unpaired two-tailed *t* test. *, $P < 0.05$; **, $P < 0.01$.
